# A robust negative association between estimated tumour circadian clock function and survival in early stage breast cancer

**DOI:** 10.64898/2026.03.15.711880

**Authors:** Vadim Vasilyev, Denise Vlachou, Sylvie Giacchetti, Georg A. Bjarnason, Tami A. Martino, Francis Levi, Robert Dallmann, David A Rand

## Abstract

Recent studies have established that the circadian clock influences onset, progression and therapeutic outcomes in a number of chronic conditions including cardio-metabolic diseases and cancer. For the latter, they also suggest that chronotherapy offers the potential to refine current treatments and improve the development of future anti-cancer medicines. Therefore, there is a need for tools to measure the functional state of the tumoural circadian clock in patients. We have previously developed a model-led machine-learning algorithm called TimeTeller which has the potential to provide such a tool. Here we demonstrate its potential for clinical relevance by a study of 1286 breast cancer patients in which we characterise the nature of the circadian clock disruption in their tumours and demonstrate a strong nonlinear association between 10-year survival and TimeTeller’s tumoural clock disfunction score Θ. We find that good tumour clock function is antagonistic to survival.

## Introduction

Breast cancer remains the most common cancer among women worldwide, with approximately 2.3 million new cases and more than 700,000 deaths reported in 2022 alone [1]. Importantly, women working night shifts experience an increased risk of developing breast cancer [2–4], contributing to the World Health Organization’s classification of circadian-disrupting shift work as a probable human carcinogen [5]. Experimental studies further show that chronic circadian misalignment accelerates tumour initiation and growth, and disrupts metabolic and immune homeostasis [6, 7].

For the treatment of patients with early stage primary breast cancer, decisions regarding adjuvant or neo-adjuvant chemotherapy or radiotherapy depending on a complex interplay of prognostic factors. These include hormone receptor status, lymph-node involvement, and, increasingly, molecular subtypes derived from gene-expression profiling [8]. Building on these molecular markers, numerous publicly available transcriptomic datasets have been extensively analysed to uncover novel gene-expression signatures associated with recurrence risk and survival [9–11]. Together, these developments have transformed patient stratification and enabled more personalised therapeutic decision-making. Yet, the lack of an appropriate tool to measure tumour circadian clock functionality and timing has prevented the integration of temporal information into patient stratification and treatment personalization attempts.

A large body of work has documented the relevance of circadian rhythms for cancer biology and treatment responses. Circadian rhythms are endogenous oscillations with a ≈24-h period and are found in most physiological parameters including rest-activity, body temperature, hormonal secretions, cellular metabolism, proliferation, death, and immune functions. These rhythms are coordinated by the suprachiasmatic nucleus (SCN), which synchronises molecular clocks residing within each cell via rhythmic signals, forming a hierarchical circadian timing system (CTS) [12]. Mammalian cells contain molecular clocks involving at least 15 clock genes (*ARNTL, CLOCK, NPAS2, PER1, PER2, PER3, CRY1, CRY2, NR1D1, NR1D2, RORA, BHLHE40, BHLHE41, DBP,* and *TIMELESS*) that interact through three transcription/post-transcription regulatory loops, generating circadian oscillations [13].

Many critical questions remain unanswered: Once cancer develops, do tumours retain functional circadian clocks? If so, how accurately do these tumour clocks keep time? And does tumour clock timing accuracy affect patient outcomes?

Previous studies suggest that malignant transformation often leads to circadian dysregulation in the cancer cells [14–17]. While some cancer cell lines exhibit robust clock function [18, 19], others appear largely arrhythmic or “clockless” [20, 21]. Altered expression of selected clock genes in tumours has been associated with more aggressive tumour phenotypes and poorer patient survival [22–26]. Yet, it remains unclear whether such misexpression of individual clock genes in tumours is a direct marker of tumour clock dysfunction.

Determining whether tumour circadian clocks help or harm patients has been hampered by the lack of tools to assess clock function in individual patient samples. To be clinically useful, such tools must evaluate individual tumour samples independently, i.e. one patient’s assessment should not depend on others, though the tool may need to be trained on separate healthy data. Moreover, they should be grounded in a statistically rational framework that allows for quantitative assessment and proper analysis of uncertainty. Existing approaches [14, 17, 27–37] estimate circadian phase but do not quantify clock functionality at the single-sample level.

To address this need, we developed TimeTeller, a model-led machine-learning algorithm designed to estimate, from a single transcriptomic sample, both the internal circadian phase (ICP), which reflects when the molecular clock assumes it to be, independent of actual wall-clock time, and the degree of clock dysfunction [38]. TimeTeller’s estimate of the ICP is called *TT time*. In [38], TimeTeller was validated using diverse murine datasets, including genetically and environmentally perturbed models, and tested on a limited set of primate and human data. TimeTeller can be used to determine whether cells have coherent clocks, while its dysfunction score, Θ, quantifies timing accuracy and oscillatory quality: a higher dysfunction score indicates a disrupted or inaccurate clock, whereas a lower value reflects a more functional clock. Examination of each sample’s timing-likelihood curve further reveals the detailed structure of the circadian signal in the tissue.

Here, we analyse molecular clock timing and functionality in primary breast cancer samples from 1,286 patients registered in one of six clinical cohorts. First, we show that most breast tumours retain mechanistically functional molecular clocks, as indicated by coherent phase predictions, gene states, and gene-gene correlations across clock genes. However, though the ICP in the tumours and the clock gene state is compatible with times in the working day, the ICP in the tumours appears to be disconnected from the daily clock hours, a finding consistent with external and possibly internal desynchronisation. Second, we identify a strong association between TimeTeller’s tumour dysfunction score and patients’ survival outcomes across multiple endpoints, including 10-year overall survival. Strikingly, we show that a well functioning tumour clock may confer a significant survival disadvantage to the patient. This second finding is consistent with recent work [26], using different methods, which indicated decreased 5-year survival for luminal A patients with a high level of a proposed measure of rhythm amplitude compared to a medium level. We further examine how these results vary with key covariates such as tumour size, lymph-node involvement, hormone-receptor and HER2 status, PAM50 subtype, and treatment.

Combining these two results, it appears that in all but a small minority of patients the tumour clock has been reconfigured so that it actively influences tumour behaviour so as to support cancer progression. We provide evidence that this is not through an increase in proliferation and suggest other mechanisms. We further show that stratifying patients by Θ revealed biologically meaningful differences in gene expression patterns and functional domains.

## Results

We analysed a cohort of 1,286 breast cancer patients from six clinical studies, namely REMAGUS (N = 226 patients), Mainz (N = 200), Rotterdam (N = 286), Transbig (N = 198), Unt (N = 125) and Upp (N = 251). We considered three survival endpoints: Overall Survival (OS), Distant Metastatic-Free Survival (DMFS) and Recurrence-Free Survival (RFS), whose availability varied among the six studies (SI Table S1), so that the final sample sizes were N = 442 patients for OS (REMAGUS and Transbig), N = 809 patients for DMFS (Mainz, Rotterdam, Transbig and Unt), and N = 776 patients for RFS (REMAGUS, Transbig, Unt and Upp).

Patient characteristics are described in detail in SI Tables S2 - S4. Briefly, the majority of the 1286 patients had small node-negative tumours, and received no systematic treatment. In contrast, the patients in the REMAGUS study included patients with locally advanced tumours who also received neoadjuvant chemotherapy with or without celecoxib or trastuzumab according to HER2 status, after the initial transcriptome biopsy was obtained. There is therefore significant heterogeneity in prognostic factors and treatment among the six studies. We regarded such heterogeneity as an opportunity to investigate possible differences in the impact of tumour circadian clock disruption on survival outcomes.

All these biopsies were analysed using either the Affymetrix Human Genome U133A Array or the expanded Affymetrix Human Genome U133 Plus 2.0 Array. For the TimeTeller analysis we used ten common clock gene probes, as detailed in SI Table S17. For the details of the data sets, microarray processing, initial data analysis and TimeTeller tuning see SI Notes S1 - S4.

As in (4), we refer to the set of probes used as the rhythmic expression profile and the corresponding vector of expression levels as the Rhythmic Expression Vector (REV).

**Key Terms Glossary**

**REV/nREV (***g*): Rhythmic Expression Vector/normalised REV – the 10-gene clock gene expression profile used by TimeTeller and the intergene normalised version

**timing likelihood curve (***L_g_*(*t*)): estimated proportional to the probability that the sample was taken at time *t* assuming that it comes from the training clock

**ICP:** internal circadian phase

**TT time (***T* ): TimeTeller-estimated ICP for a tumour sample

**sample time (***t_s_*): Actual wall clock time when the biopsy was collected

*g*_1_: Projection of REV onto first principal component

**ML:** Maximum likelihood value of the timing likelihood curve

### Training on healthy clocks enables measurement of tumour clock deviation

TimeTeller learns from training data taken from healthy cells the quantitative structure of both the temporal variation in the expression of the genes in the REV and in the correlations between them (SI Figs. S1, S2). It does this by analysing the training data time-series. After training, this learnt probability model can be used to analyse independent test samples to estimate the likelihood that the sample was collected at any given time of day. The resulting estimated *likelihood function L_g_*(*t*) is defined for all times *t* around the day and can be regarded as proportional to the probability that the sample was taken at time *t* according to the above learnt probability model. It depends on the sample’s REV *g* and the training data, but is independent of any other such test sample and therefore can be independently estimated from a single patient sample.

Using this likelihood function, TimeTeller provides three clinically relevant summary measurements from a single tumour biopsy. First, the tumour’s ICP (TT time), which reflects when the tumour’s molecular clock assumes it to be, is defined as the time point at which the likelihood function reaches its maximum value (denoted ML), representing the most probable circadian phase. Second, the maximum likelihood value ML itself, which measures how closely the REV resembles a high-probability sample from the training clock. Third, and critically, a *dysfunction score* Θ for the sample that measures how well the tumour clock is functioning compared to the clock in healthy tissue by integrating multiple features of the likelihood curve: the sharpness of curvature near the peak, the maximum likelihood (ML) value, and the presence of prominent secondary peaks.

Examples of likelihood functions from patient samples are shown in Fig. 1A. A key observation is that likelihood curves for tumour samples are markedly more heterogeneous than those from healthy control tissues (Figs. 1A,B). To make the shapes more visible, curves have been temporally translated so their maxima align at noon and scaled by their maximum value ML to normalize the peak at 1.

**Figure 1:**
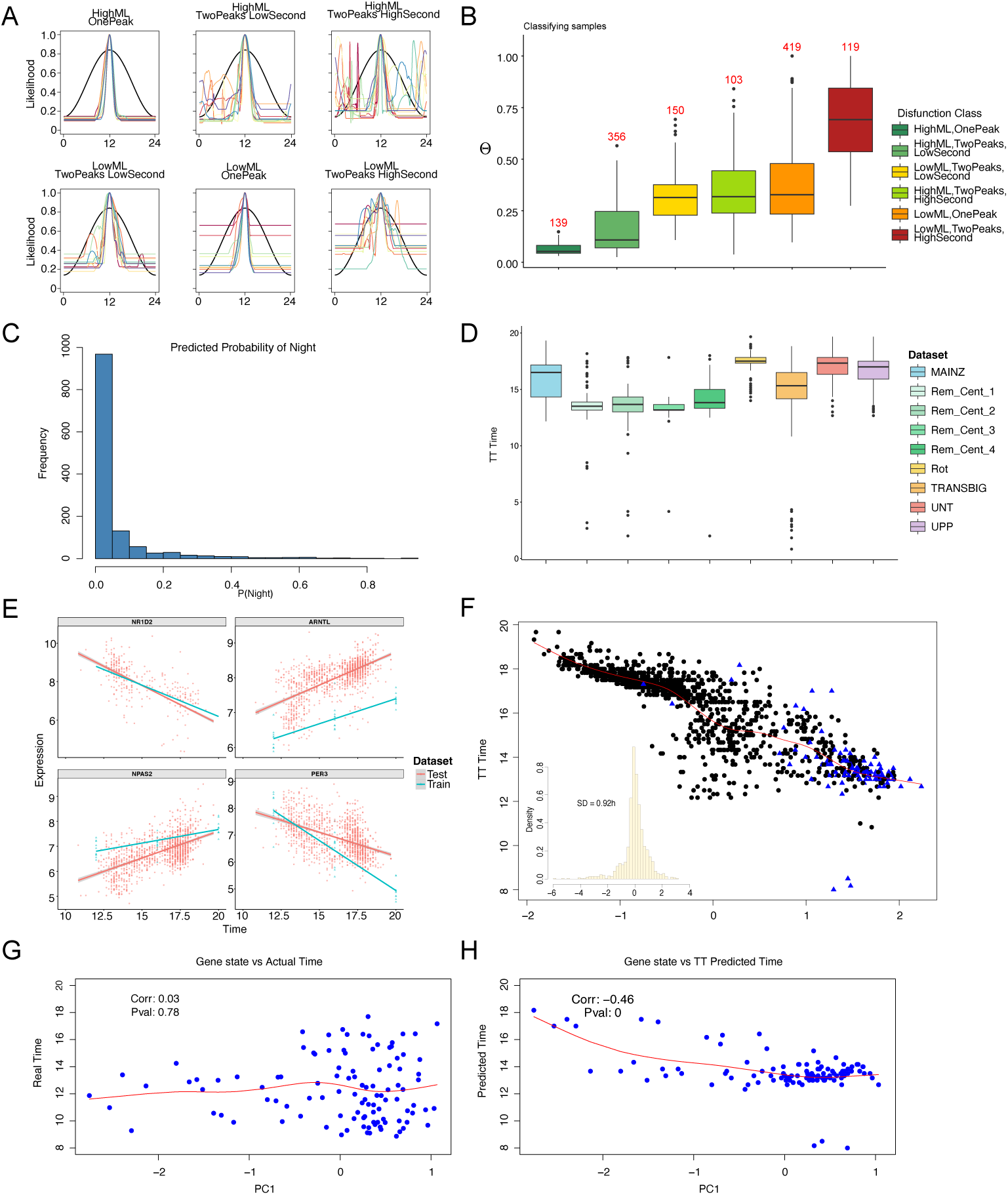
Breast tumour transcriptomes retain coherent circadian timing information despite temporal desynchronization. **A**. Representative TimeTeller likelihood curves illustrating six distinct dysfunction patterns observed across the cohort. Curves are normalized to unit maximum and temporally aligned with peak at 12:00 for visualization. The six groups shown are sampled from the six dysfunction classes shown in B. **B**. Distribution of dysfunction scores (Θ) stratified by likelihood curve morphology (corresponding to patterns in **A**). Categories are defined based on maximum likelihood (ML) value and peak structure: High ML/single peak (healthy-like), High ML/two peaks with low or high secondary peak, Low ML/single peak, Low ML/two peaks with low or high secondary peak. **C**. For each sample we use the LASSO classifier to estimate the probability that a sample is from the night. Histogram of results for the breast cancer samples. **D**. Distribution of TT time estimates by study center, demonstrating inter-center variability in phase distribution while maintaining consistent clock gene expression patterns (see panel **E**). **E**. Clock gene expression profiles as a function of TT time for four representative genes (*NPAS2*, *PER3*, *NR1D2*, *ARNTL*). Triangles indicate training data (healthy tissue); solid line shows fitted linear regression. Gene expression shows coherent circadian variation consistent with training clock despite tumour context. **F**. Relationship between TT time and value (*g*1) of projection onto the first principal component of the ten-probe normalized rhythmic expression vector (nREV), across all samples. Red curve: smoothing spline estimating *E*[*T* |*g*1] (correlation *r* = −0.34, *p <* 2.2 × 10*^−^*^16^). This tight relationship enables estimation of conditional distribution *P* (*T* |*g*1). *Inset:* Conditional distribution *P* (*T* |*g*1) showing temporal precision of TT time estimates given gene state (SD ≈ 0.92 h). **G**. External sampling time versus *g*1 for REMAGUS cohort samples with recorded collection times (N = 108). Absence of correlation (r = 0.03, p = 0.78) indicates temporal desynchronization between tumour clock state and external time. **H**. TT time versus *g*1 for the same 108 timed samples, demonstrating preserved strong correlation (r = −0.46, p = 0) between internal tumour clock phase and gene expression state despite loss of external synchronization shown in H.

The shape of the likelihood curve reveals critical information about tumour clock function by providing insight into different modes of circadian dysfunction (Fig. 1A,B). A sharp, single peak with high ML indicates the tumour clock closely resembles a high probability sample from the training clock and hence corresponds to a functional clock. Progressively less sharp maxima, lower ML values, or multiple competing peaks indicate increasing degrees of circadian dysfunction: flatter curves suggest uncertainty in timing; lower ML indicates weaker match to the training clock; multiple peaks reveal conflicting circadian phases. Fig. 1A illustrates six representative dysfunction patterns observed across our cohort, ranging from healthy-like (sharp, high peak) to severely disrupted (flat or multi-peaked curves). The dysfunction score Θ ranges from 0 to 1. Values near 0 correspond to sharp, unimodal likelihood curves with high ML, indicating well-functioning circadian clocks similar to healthy tissue (which typically exhibits Θ *<* 0.1), while values approaching 1 indicate flat, multi-peaked, or low-ML curves reflecting severe circadian dysfunction. Fig. 1B shows the distribution of Θ values stratified by likelihood curve morphology, demonstrating that curve patterns correspond systematically to dysfunction severity. In our cohort, of early stage breast cancer patients, approximately 50% of tumours showed high-confidence functional clocks (low Θ, high ML), ∼35% showed moderate dysfunction, and fewer than 15% exhibited severely disrupted clock signatures. Importantly, TimeTeller quantifies these dysfunction modes on a per-sample basis, enabling individualized tumour clock assessment not possible with population-level methods. This is discussed in further detail in [38].

### Breast tumour clocks remain mechanistically functional

Determining whether tumour circadian clocks remain mechanistically functional is important because, if so, they may actively influence tumour behaviour, either promoting or suppressing progression. By mechanistically functional we mean that they continue as a functioning molecular machine to diurnally oscillate with significant coherent clock gene expression and appropriate gene-gene correlations. It does not necessarily mean that the clock behaves or is regulated like the clock in healthy breast cells and, at this stage we are not implying positive or negative clinical effects. We therefore used TimeTeller to systematically evaluate this for the 1,286 breast tumour samples.

We investigated the TT time for each sample and found that for all but 88 (6.8%) of the 1,286 tumour samples, the estimated TT time fell within the working day window which we take as 7am to 8pm. We examined the likelihood curves for the samples yielding off-target TT times and found that 66 (76%) of them exhibited a secondary peak within the working day window. For these samples, we reassigned TT time to the secondary peak. This adjustment is biologically reasonable given that likelihood curves with multiple peaks indicate competing times. The remaining 22 off-target samples (1.6% of total) maintained their original TT time assignments.

We were surprised that the TT times for these breast cancer samples fell overwhelmingly within working hours because, based on previous analysis of cancer data [26, 39], we expected to see a more chaotic broader spread of times. We therefore decided to do a separate analysis independent of TimeTeller to determine whether the observed clock gene states were compatible with daytime or nighttime. To do this we studied the ratios of expression of pairs of probes in the REV and compared these with what would be expected in the training clock. We used lasso logistic regression to identify the minimal set of gene expression ratios capable of discriminating day from night phases (SI Note S5). The final model selected 5 pairwise ratio features (SI Fig. S18) and achieved 97% classification accuracy on the training data, indicating that a sparse subset of clock gene relationships is sufficient to capture day/night phase information. When applied to our data only 24 of the 1286 samples are classified during nighttime and nearly a thousand have probability of more than 95% of belonging to daytime (SI Fig. S18) supporting the TimeTeller analysis.

The observation that TT times fall predominantly within working hours despite samples being analyzed independently, provides initial evidence that tumour clocks maintain approximately 24-hour rhythms. We validated this finding and assessed clock functionality through multiple complementary approaches.

First, we checked for batch effects between centres. The TT times for the individual centres showed significant variation between centres (Fig. 1D, Figs. S3,S8a). However, when the data from the centres were combined and the expression levels of the REV genes were plotted against TT time, the gene expressions changed in a coherent roughly linear way with a clear positive or negative slope for the genes that vary across the daytime and were flat otherwise (Fig. 1E, SI Figs. S4, S5, S6).

This result suggested that for the REVs any batch effects between the centres were small and the observed differences in the timing between centres were real. This was further supported by PCA plots (SI Fig. S7). We also observed that this variation in clock gene expression agrees with the expression levels of the training data across this period (Fig. 1E).

Next we investigated whether we could extend this relationship to a coherent near-linear relationship between TT time and the multidimensional gene state as encoded by the nREV. Moving on from individual genes to the nREVs is important as the nREVs contain much more information about the internal phase and its variability and they encode the temporally changing relationships between the REV genes.

To determine the correspondence between TT Time and REV we utilised the projection *g*_1_ of the nREV *g* onto the first principal component (PC) of the set of all 1286 nREVs (Fig. 1F). Since we observed that the TT times of the samples were almost all in the working day and the variation of the REV genes was approximately linear over these times (Fig. 1E and SI Figs. S4 & S5) using PCA to approximate the data is reasonable. Indeed, we found that the first PC captured a large proportion (just over 40%) of the relevant variance in the REV because of this.

Plotting TT time against the first PC (Fig. 1F) showed very good correlation between them (slope −0.86, p-value *<* 2.2e-16). Moreover, the TT time was tightly constrained by the first PC *g*_1_ as, using all 1286 samples, we found that, conditional on knowing *g*_1_, the TT time *T* had a standard deviation of approximately 1h (Fig. 1G). This accuracy compares very favourably with the typical standard deviation between estimated time and predicted time in previous studies of human tissue data which typically have a median absolute error greater than 1.4h [17, 31, 32]. We concluded that the TT time was well determined by the REV.

It follows that TT time is given as a near linear function of the nREV with an error having a standard deviation of an hour or less. This result is strengthened by the observation that if we used the value of ML to split the data into four quartiles there was a significant increase in concordance between TT time and *g*_1_ as the ML value increases (SI Fig. S8b). A high value of ML relative to the values obtained by training data is indicative of the fact that the REV comes from a clock that is similar to the training clock. In particular, we noted that for the top quartile, the standard deviation of the TT times conditional on knowing *g*_1_, had shrunk from approximately 1h to 0.7h (SI Fig. S12a,b). Strikingly, for the half of the patients with highest ML, the TT time well determines the REV in that the SD of *g*_1_ conditional on knowing the TT time at 0.37 is approximately 10% of the spread of *g*_1_ across the working day for the patients with the top half of ML values and at 0.54 just approximately 15% for the lower half (SI Fig. S13). This means that up to addition of moderate noise the gene state is given by the TT time, providing strong evidence of a working oscillator.

Previous work on untimed breast cancer samples [26] has confirmed that clocks are oscillating and not stalled using different methods based on the population of patients rather than individual patients. In particular, checking the population-averaged gene-gene correlations for core clock genes across the population reflects the behaviour of physiological clocks and/or by fitting a nonlinear regression to clock gene expression vs estimated time and testing that this shows circadian variation. We checked that similar population analyses of our data further validated our conclusion that the breast cancer tumour circadian clocks are mechanistically functional (SI Note S6, SI Fig. S17).

In summary, these results provide strong evidence for diurnal oscillations in tumours because it shows that given the REV gene state for a single patient’s tissue sample we can systematically predict a patient specific phase using TT time with a SD error of about an hour (and better if the ML is high) and, moreover, the result for such a patient falls in a consistent pattern with the other patients. The coherence that we consistently find in clock gene expression as a function of TT time provides strong evidence that tumour samples occupy different circadian phases in a manner consistent with functional oscillators. Rather than random or frozen gene expression patterns, tumours show the systematic phase-dependent variation expected from working oscillators.

### Tumour clocks lose external synchronisation

Having established mechanistic functionality with TT times in the working day and that TT time accurately reflected tumour clock gene state, we next asked whether tumour clocks maintain synchronisation with external time. Only 108 samples had sampling time available, all from the REMAGUS trial. In these samples, sampling time was not correlated with either TT time (slope −0.027, p = 0.80) or gene state *g*_1_ (Fig. 1H), contrasting sharply with the tight correlation between TT time and *g*_1_ (Fig. 1I).

In healthy human tissue, some deviation from sampling time is expected due to chronotype variation (i.e., earlier or later diurnal preference), which is partly genetically based [40]. However, if this lack of correlation were due to chronotype, differences between TT time and sampling time should be approximately normally distributed with standard deviation ≈1 hour. Instead, the observed distribution is far from normal with much larger standard deviation (Fig. S9).

This complete lack of correlation between external sampling time and internal tumour time represents fundamental disruption in the circadian timing system. Importantly, this is not clock absence but temporal uncoupling: tumour clocks are running but have lost synchronization with the external light-dark cycle and central pacemaker. This temporal desynchronization could reflect loss of SCN control, tumour microenvironmental effects that override entrainment signals, or active selection for autonomous clock function, mechanistic disruptions that may be therapeutically targetable.

Interestingly, tumour circadian phases (i.e., TT times) are not uniformly distributed but show temporal compression, clustering rather than spanning the full working day. This is clear for samples with known timing (Figs. 1C,D), which cluster around midday, and also indirectly from tight clustering of times within each centre (SI Fig. S3).

We conclude that although the clock is mechanistically functional in most samples, with gene state tightly constrained by TT time and phases consistently falling in the daytime, TT phases and gene states are incompatible with sampling time when known. Thus, the tumour clock has been substantially modified.

### Patient survival is strongly linked to dysfunction score Θ in primary breast cancer tumours

Given that tumour clocks are mechanistically functional but fundamentally altered, and that host circadian disruption has been implicated in cancer progression [2, 3, 5–7, 41–47], these findings raise a critical question: What is the clinical significance of better versus worse clock function? Does preserved clock function benefit patients by maintaining cellular organization, or harm them by enabling tumours to exploit circadian coordination? We therefore examined the association between TimeTeller’s dysfunction score Θ and clinical outcomes.

Due to differences in knowledge and diagnostic methods at the time the six studies of interest were conducted, the reported prognostic factors (including lymph node status, tumour size or histological grade, and hormone receptor and HER2 status) differed between the studies, thus preventing the incorporation of a universal set of covariates across all centres. To partially overcome this limitation, we incorporated the PAM50 molecular classification [48], which retrospectively stratified the 1286 breast cancers into five intrinsic subtypes (Luminal A, Luminal B, HER2-enriched, Basal-like, Normal-like) based on gene expression (for more details see SI Note S7).

As the patient characteristics varied significantly between centres, we decided to consider the data from the REMAGUS trial first, as this represented a relatively homogeneous group of patients with both a full report on the classical prognostic factors and comparatively large tumours, all treated with chemotherapy. This multicenter randomised phase II clinical trial aimed to assess the response of primary breast cancer to different protocols of neoadjuvant chemotherapy according to tumour hormonal receptor status and HER2 expression [49–52]. Of the trial’s 340 patients, 226 had a pretreatment cancer biopsy using the same RNA extraction procedure and analysed with Affymetrix Human Genome U133 Plus 2.0 Arrays.

Kaplan-Meier survival analysis of the dysfunction score Θ requires the choice of a threshold since Θ is a continuous variable. As the training data is meant to represent tissues with a good clock function, a robust and meaningful threshold is the value of Θ at the start of the upper 90th percentile of the training data. We designated samples below this threshold as having good clock function (the “good clock” group, GCG) and those above as having bad clock function (the “bad clock” group, BCG). With this choice the analysis produced the overall survival curves shown in Fig. 2A, revealing a statistically significant survival advantage for the “bad clock” patients.

**Figure 2:**
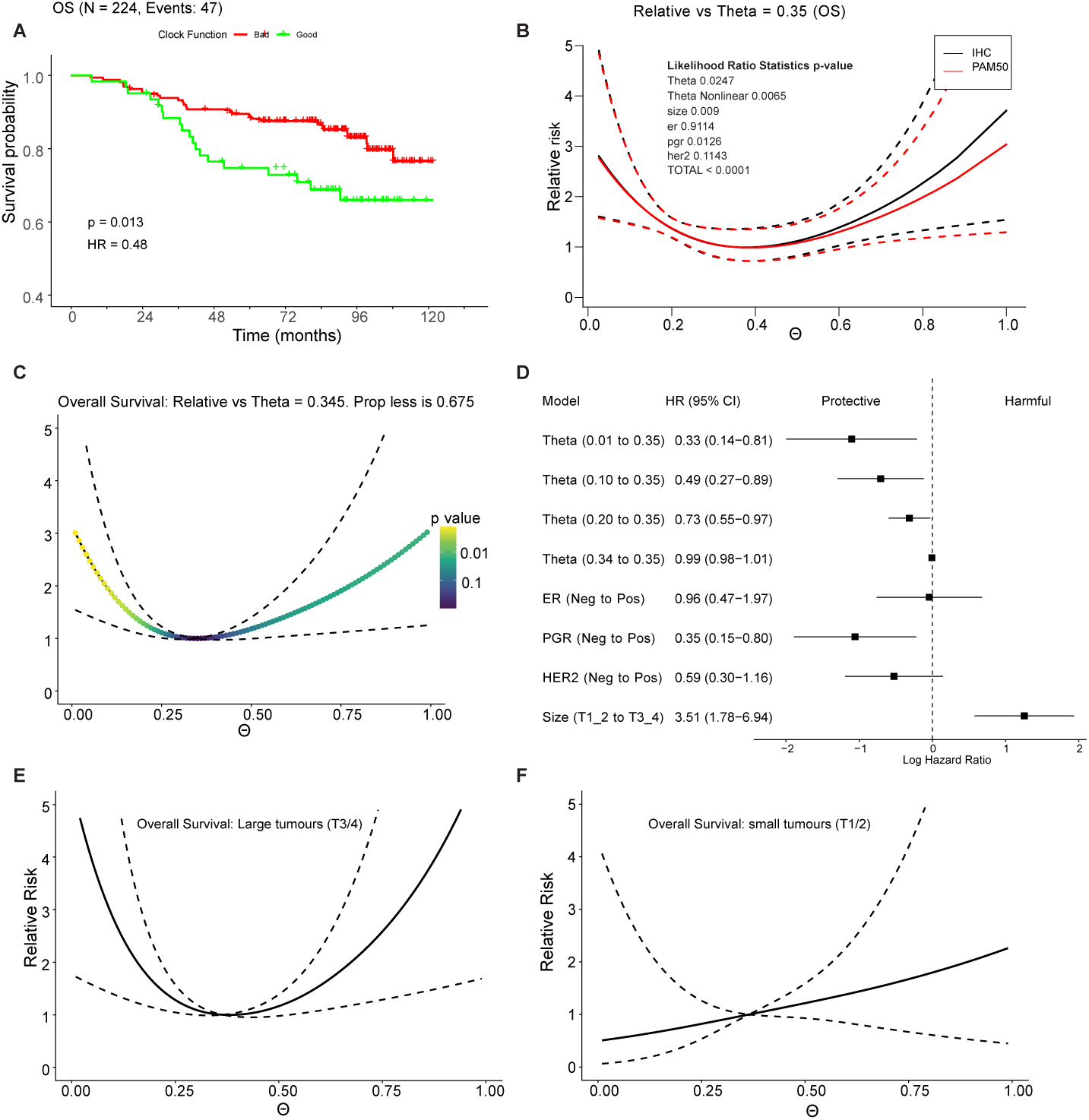
Strong nonlinear relationship between circadian dysfunction score Θ and overall survival in REMAGUS breast cancer patients. All models use restricted cubic splines with knots at Θ quantiles (default) to capture nonlinearity. **A**. Kaplan-Meier survival plot showing differences in overall survival (OS) for patients above and below the critical Θ value defining the good and bad clock groups. (HR = 0.48, log-rank *p* = 0.013). **B**. Relative risk as a function of Θ for the REMAGUS patients with Θ = 0.35 as reference point (relative risk = 1). Black curve: model including ER, PR, HER2 status as covariates; red curve: model substituting PAM50 molecular subtype as covariates. Both models adjust for age and tumour size (T1–T4). Approximately 80% of patients have Θ *<* 0.4. Risk decreases substantially from Θ = 0 to Θ ≈ 0.35 (3-fold reduction), then plateaus. Dotted lines show 95% confidence intervals around the estimated relative risk curve. Inset: likelihood ratio test statistics demonstrating highly significant nonlinearity (*p* = 0.0009), Θ effect (*p* = 0.003), and tumour size effect (*p* = 0.0001). Confidence intervals around the relative risk estimate are shown. **C,D**. Panels C–E show confidence intervals for the contrast (difference) between a given Θ and Θ = 0.345; these necessarily converge at Θ = 0.345. **C**. Relative risk as a function of Θ for all the patients with statistical significance heatmap showing *p*-values for pairwise comparisons of relative risk at arbitrary Θ values versus the risk-minimizing value (Θ = 0.345). Highly significant differences observed for Θ *<* 0.25, confirming that low Θ (better clock function) associates with substantially worse outcomes. Dotted lines indicate 95% confidence intervals for the contrast, which converge at Θ = 0.345 where the contrast is zero by definition. **D**. Forest plot showing hazard ratios and confidence intervals for Θ and other covariates from multivariate analysis using nonlinear Cox model. **E, F**. Relative risk stratified by tumour size through inclusion of Θ×size interaction term, showing contrast between given Θ values and reference Θ = 0.345. **E**. Large tumours (T3/T4) show dramatic risk reduction (nearly 5-fold) as Θ increases from 0 to 0.345. **F**. Small tumours (T1/T2) show modest, nearly flat relationship between Θ and risk, indicating the protective effect of circadian dysfunction is predominantly observed in advanced-stage disease. In both panels, 95% confidence intervals (dotted lines) converge at Θ = 0.345 where the contrast equals zero.

This finding warranted careful interpretation in view of the reported association of tumour clock disruption and cancer progression and worse outcomes (e.g. [14, 15, 22, 47, 53]). Moreover, a similar analysis splitting the population into a “bad”, “average” and “good” clock group using a 25:50:25 split gave a result where the “average” group had a significant survival advantage over the other groups (Fig. S10a) raising the possibility of a nonlinear relationship between dysfunction and survival. These unexpected results led us to investigate the relationship more deeply using a more powerful analysis treating Θ as a continuous variable rather than using a binary threshold.

The usual choice for such an analysis is the Cox proportional hazards model [54]. This model typically assumes a linear relationship between risk factors (like Θ) and survival. However, when there is evidence of nonlinearity, as we suspected for Θ, restricted cubic spline functions can be incorporated to detect and characterize these nonlinear effects [54] (SI Note S8 for more details).

The nonlinear multivariate model produced a striking, statistically significant relative risk curve for the REMAGUS OS data (Fig. 2B,C). In this case, the relative risk curve gives the hazard ratio of the risk of earlier death at a given value of Θ relative to that at the baseline value (here Θ = 0.35). Three key findings emerged. Firstly, both the non-trivial dependence of relative risk on Θ and the presence of nonlinearity were highly significant (resp. *p <* 0.003 and *p <* 0.002). Secondly, relative risk decreased substantially as Θ was increased from 0 to approximately 0.35, a Θ range that included nearly 70% of patients’ tumours. At Θ = 0.35, the relative risk was reduced by nearly threefold. Thirdly, the relative risk then increased for Θ *>* 0.35, but with wide confidence limits for Θ in this region (Fig. 2C). These results held whether we used hormone receptor status (ER, PR, HER2) or PAM50 molecular subtypes as covariates (Fig. 2B).

The significance of the Θ-dependent survival, the presence of nonlinearity, and the possibility of interactions between Θ and other variables was assessed using a likelihood ratio test ( [54] and SI Sects. S8). Statistics of all model tests are in SI Tables S5 - S16. We also analysed the data for Θ *<* 0.35 and Θ *>* 0.35 separately and showed that for tumours with Θ *<* 0.35, the relative risk curve decreased monotonically, and the results for Θ became even more significant (*p* = 0.009), whereas the result for patients with Θ *>* 0.35 was not significant.

We considered whether the change in the sign of the slope of the relative risk curve might have been due to an interaction with either tumour size or cellular proliferation rate, since we saw from our results that changes in either could have a strong effect on survival. To test the effect of size, we added an interaction term between Θ and size to the model (SI Note S8). This revealed a striking difference: while patients with small tumours showed a modest decrease in relative risk as Θ increased, patients with large tumours showed a dramatic drop, with relative risk falling by nearly 5-fold over the range 0 *<* Θ *<* 0.35 (Fig. 2D,E).

Clinically, this finding about the interaction could inform risk stratification because it strongly suggests that large tumours with well-functioning clocks (low Θ) may be more capable of sustaining aggressive growth and, though less convincingly, that the same is true for large Θ.

Similar results were obtained for OS when the other patients with OS data were added to those from the REMAGUS dataset (N = 422, Fig. 3A-C). However, unlike the REMAGUS patients, the added patients (N = 198, from the Transbig centre) did not receive treatment and this enabled us to include treatment as a covariate. For the full OS group there was a highly significant difference, with treated patients showing substantially better outcomes than untreated ones (HR = 0.59, p=0.016, Fig. 3A). Critically, the association of the dysfunction score Θ with survival remained highly statistically significant even after adjusting for treatment status. This indicates that Θ provides independent prognostic information in that it predicts outcomes regardless of whether patients receive chemotherapy or not. This independence from treatment suggests that Θ reflects intrinsic tumour biology rather than treatment response, and could potentially be used to stratify patients in both treated and untreated populations.

**Figure 3:**
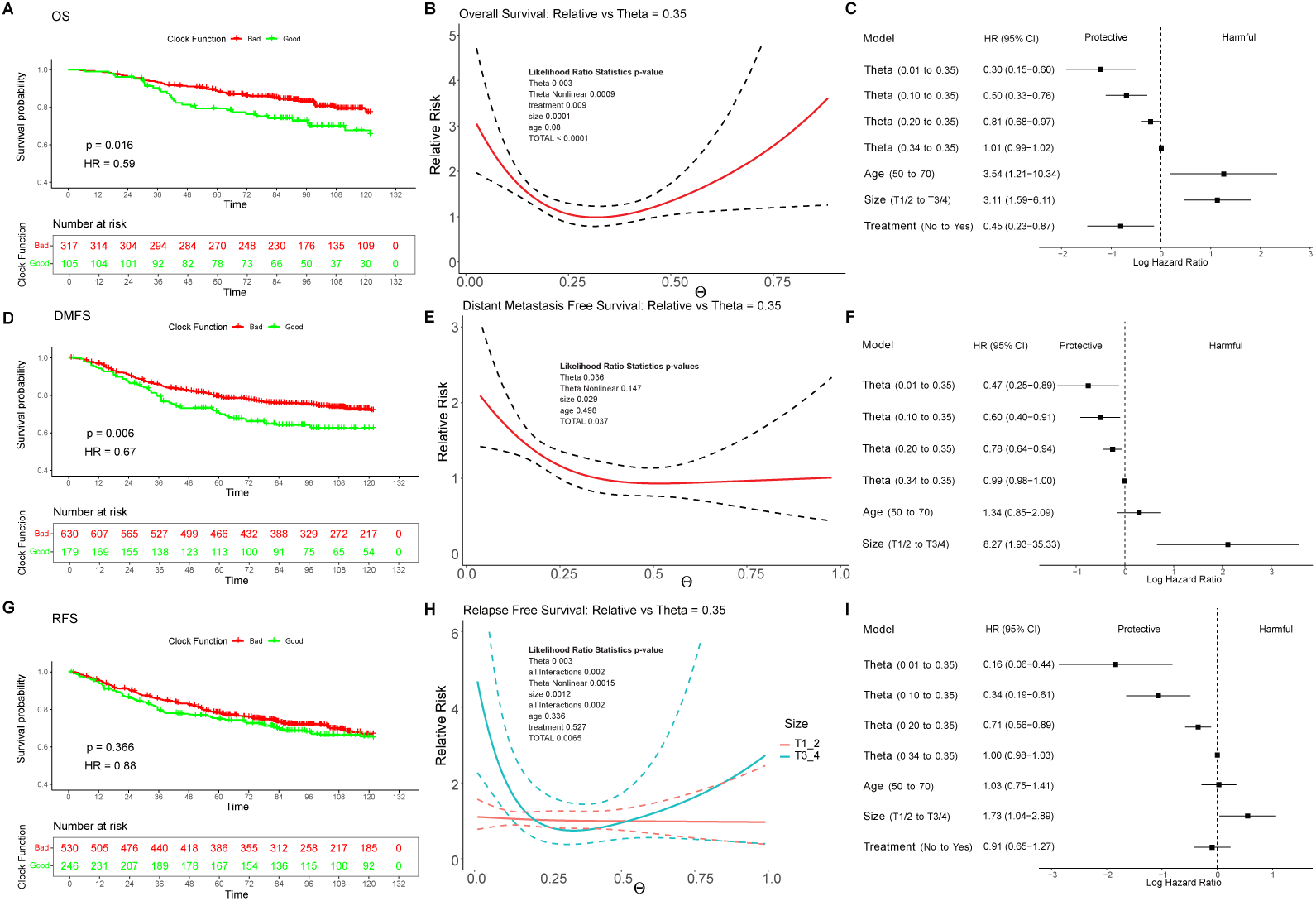
Survival results for OS, DMFS and RFS. **A**. Kaplan-Meier survival plot for patients in the full OS group showing differences in overall survival (OS) above and below the critical Θ value defining the GCG and BCG. **B**. Relative risk as a function of Θ for the patients in the full OS group calculated with the nonlinear Cox model. The p-values from the likelihood ratio analysis are shown with meaning as in Fig. 2. **C**. Multivariate analysis of the patients in the full OS group. As well as the items listed PAM50 status is included as covariates. **D-F**. As A-C but for the DMFS group. **G,I**. As A-C but for the RFS group. **H**. As B except for the RFS group and since an interaction between Θ and size is shown to be statistically significant for this group, a corresponding interaction term is included and the separate plots for small:T1/2 and large:T3/4 are included.

For the DMFS group, the questions about size and treatment was irrelevant since there were no patients in the large tumour group and none were treated. While in this case the nonlinearity was not statistically significant we included it in line with recommended practice [54]. The Θ dependence and size were statistically significant and the relative risk as a function of Θ was monotonically decreasing with approximately a halving of risk by the time Θ = 0.5 (Fig. 3D-F).

For RFS the presence of the interaction term between Θ and size was statistically significant and therefore had to be included. As for OS, the analysis supported the hypothesis that there might be an interaction between size and Θ, and provided evidence of a more than fourfold drop in relative risk for patients with larger tumours (Fig. 3G-I). Since the slope of the curve for small tumours was almost flat (Fig. 3H) it also supported the idea that the interaction caused the change in the sign of the slope (Fig. S10c).

To investigate whether the association between tumour clock functionality and survival outcomes involved clock-controlled proliferation pathways, we constructed an index using established proliferation markers (SI Note S9). The Pearson correlation between Θ and this proliferation index and Y was negligible. Although the Pearson correlation at 0.07 was statistically significant (*p* = 0.02, 95% CI [0.01, 0.13], *n* = 1,050), the effect size was negligible, with the correlation explaining less than 0.5% of the variance (*R*^2^ = 0.005). This suggests no meaningful linear association between the two variables supporting tumour circadian dysfunction as an independent and new prognostic indicator. To further understand how the effect of Θ varies across breast cancer subtypes, we separately analysed each PAM50 subgroup (SI Figs. S21 - S24) and found that the effect of Θ on overall survival varies substantially by molecular subtype. For the Basal subtype which is predominantly triple-negative there was a significant association between higher Θ and improved survival (p = 0.02). Given that Basal/triple-negative tumours typically have poor prognosis and limited treatment options, Θ may be particularly valuable for risk stratification in this subgroup. On the other hand, for the other three subtypes the effects were smaller and non-significant but the trend was consistent.

This may reflect better baseline prognosis, different biology and is also affected by the smaller sample sizes. These subtype-specific effects suggest that circadian dysfunction interacts with the underlying molecular drivers of each breast cancer subtype, and that Θ may be most clinically useful in aggressive subtypes.

In summary, we identified a strong, statistically significant association between tumour circadian dysfunction (Θ) and patient outcomes. For the 70-80% of patients with lowest Θ values (least disruption) relative risk decreases substantially with increasing Θ (more disruption). The protective effect of higher Θ was particularly pronounced in large tumours with a nearly 5-fold reduction in risk, compared to modest effects in small tumours. Moreover, we see remarkably consistent behaviour across all survival end-points and study centers despite the heterogeneity of the patients’ characteristics, lending strong support to the basic result above. The survival dependence upon Θ was independent from treatment suggesting that this dependence reflects intrinsic tumour biology rather than treatment response, and could potentially be used to stratify patients in both treated and untreated populations. The association was particularly strong in Basal (triple-negative) tumours and Luminal B tumours - precisely the subtypes with worse prognosis where improved risk stratification is most needed. The prognostic value of Θ was independent of chemotherapy status, indicating it reflects intrinsic tumour biology. There was no correlation between Θ and proliferation index, indicating Θ provides orthogonal prognostic information.

### TimeTeller’s Θ stratification uncovers biologically distinct groups

To understand how Θ relates to biological and patient characteristics, we examined whether Θ-based stratification maps onto standard biological differences such as differential gene expression or pathway activation. For this analysis, we defined two groups: a Good Clock Group (GCG) with low Θ values (better circadian function, more similar to healthy clocks) and a Bad Clock Group (BCG) with high Θ values (greater circadian dysfunction). We examined biological differences between these groups using multiple GCG/BCG definitions appropriate to the aspect being considered.

We start by considering differential gene expression. Previous work showed that stratification by TimeTeller’s dysfunction metric Θ uncovered distinct strata in oral squamous cell carcinoma biopsies [38]. We therefore tested whether similar results held for breast cancer patients.

For this analysis, we used only cohorts from the Affymetrix U133A platform (N = 1050). While clock gene expression showed no batch effects, PCA analysis (Fig. S15a) of all other common genes suggested a batch effect in the REMAGUS dataset (N = 226) compared to the rest. This could arise from biological and/or technical factors (U133 Plus2.0 array vs U133A), so we carried out separate analyses for REMAGUS and non-REMAGUS datasets, focusing here on the larger, more homogeneous non-REMAGUS cohort (Fig. S15b).

To define Good and Bad Clock Groups (GCG and BCG) objectively, we fixed an equal group size *n* for both groups, defining the GCG (resp. BCG) as the *n* tumour samples with smallest (resp. largest) Θ values. We identified differentially expressed genes (DEGs) that were statistically significant between these groups, adjusting for covariates, batch effects, sample quality and *p*-values for multiple testing (SI Note S10).

The number of DEGs between GCG and BCG increased monotonically from ≈300 genes when group size *n* was 100 to ≈1600 genes when *n* ≈ 200 (Fig. 4A, Fig. S16b), potentially due to increased sample numbers. As group size increased further to 500, DEG count reduced to a roughly constant level of ≈1200 genes.

**Figure 4:**
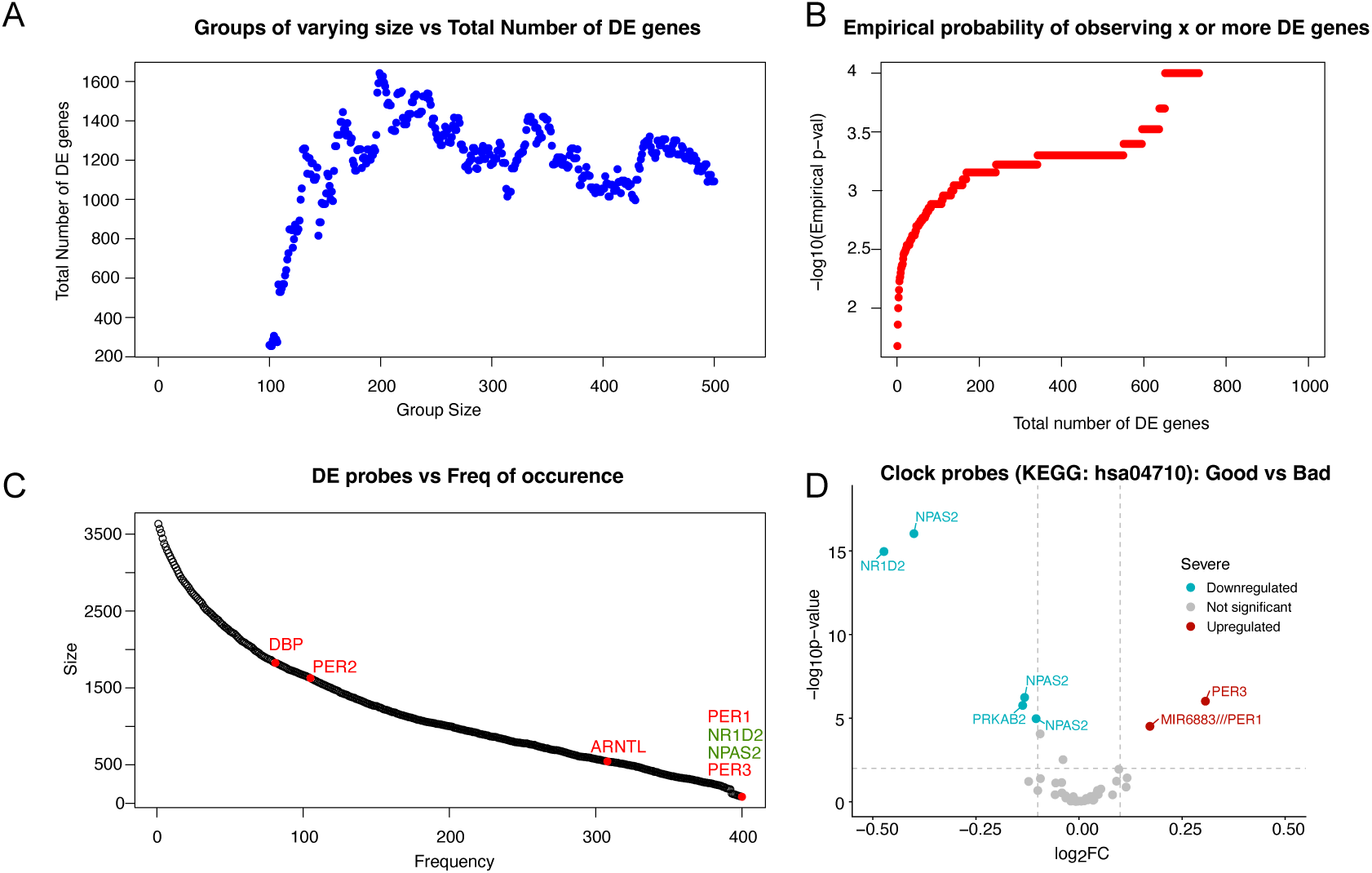
Θ-based stratification reveals distinct transcriptomic and pathway signatures between Good Clock Group (GCG) and Bad Clock Group (BCG). **A**. Number of differentially expressed genes (DEGs) as a function of equally-sized GCG and BCG group sizes (*limma* pipeline with covariate and sample quality adjustment, FDR-adjusted *p <* 0.05). Peak at *n* ≈ 250 per group yields ∼1,600 DEGs; plateau at ∼1,200 DEGs for larger group sizes as Θ distributions begin to overlap. **B**. Permutation test results showing empirical probability of observing DEG counts under null hypothesis of random clock function assignment (covariates held constant). Observed DEG counts (panel A) far exceed chance expectations (*p <* 0.001 for all group sizes tested), confirming biological significance of Θ stratification. **C**. Frequency distribution of DEG detection across varying group sizes from panel A. Approximately 100 genes show differential expression consistently across all group sizes tested, with strong enrichment for core circadian clock genes (labeled: PER1, ARNTL, NPAS2, PER3, NR1D2, DBP, PER2). **D**. Volcano plot of KEGG circadian rhythm pathway genes (hsa04710) comparing GCG versus BCG (25:50:25 split, *N* ≈ 250 per group). Multiple clock genes show significant differential expression, indicating coordinated disruption of circadian transcriptional programs in high-Θ tumours.

To test whether this pattern reflected true biological differences or random variation, we per-formed permutation testing. We randomly assigned samples to “pseudo-GCG” and “pseudo-BCG” groups of matched sizes and counted DEGs, while keeping other covariates fixed. (Fig. 4B, SI Fig. S16a). The number of DEGs observed with true Θ-based stratification far exceeds what would be expected by chance (*p <* 0.001 for each group size tested). Given that we tested multiple group sizes (*n* = 100 − 500) and all showed highly significant results, we can conclusively reject the null hypothesis that Θ stratification is unrelated to gene expression patterns.

Fig. 4C addresses gene identification consistency. Approximately 100 distinct probes showed differential expression across all group sizes, indicating these genes robustly distinguish GCG from BCG regardless of threshold. These consistently expressed genes show strong enrichment for core circadian clock genes including *DBP*, *PER2*, *ARNTL*, *PER1*, *NR1D2*, *NPAS2*, and *PER3*. Notably, *PER1*, *NR1D2*, *NPAS2*, and *PER3* were differentially expressed for *all* group sizes tested. This enrichment provides independent validation that Θ indeed captures circadian clock function. The metric was designed to measure clock dysfunction based on the most rhythmic subset of the clock genes, yet when we perform unbiased genome-wide differential expression analysis between Θ-stratified groups, we independently rediscover differential expression of circadian clock genes. This cross-validation strongly supports Θ’s biological validity.

We conclude that Θ captures real, functionally relevant biological variation in tumour transcriptomes, not simply technical noise or arbitrary variation. These differences in gene expression would not be seen without the stratification.

### Pathway analysis

To identify coordinated biological processes affected by circadian dysfunction, we performed pathway enrichment analysis comparing GCG and BCG groups (SI Note S11). For this analysis, we used a 25:50:25 split (top 25% lowest Θ vs. bottom 25% highest Θ, excluding the middle 50%) to maximize separation while maintaining adequate sample size (*N* ≈ 250 per group). Results were robust to variations in this threshold (SI Fig. S20).

Using KEGG pathway over-representation analysis (Fig. 4D), we found that the circadian rhythm pathway was significantly enriched in both up-regulated and down-regulated gene sets, indicating consistent but complex deregulation of circadian mechanisms in high Θ (BCG) tumours. This was supported by the changes seen in the volcano plot Fig. 4D which showed multiple circadian genes showing strong differential expression with high statistical significance, consistent with our earlier finding of clock gene enrichment.

Biologically, high circadian dysfunction (high Θ) associates with coordinated changes in circadian pathway gene expression, not just isolated changes in individual clock genes, suggesting system-level disruption of circadian regulation rather than random perturbations.

To further investigate the transcriptomic signatures of the groups identified by TimeTeller, we employed the Pathway RespOnsive GENes (PROGENy) algorithm, which leverages a large compendium of publicly available perturbation experiments to infer pathway activity. Rather than relying on the expression of pathway members, PROGENy focuses on downstream target genes that are consistently deregulated in response to pathway perturbation. This results in a robust set of pathway-responsive gene signatures that reflect the functional output of pathway activity [55]. This approach is particularly well-suited for distinguishing pathway activation states, as it captures the transcriptional consequences of signalling events rather than mere presence or expression of pathway components.

We first tested for associations between pathway activity and sample groups within our dataset, following the strategy outlined in Schubert et al. [55]. This enabled us to identify key signalling pathways that differentiate the TimeTeller-defined groups based on their downstream transcriptional effects. We found NF*κ*B/TNF*α* and EGFR/MAPK, which are biologically closely related and known to cross-activate to be highly correlated (Fig. 5A). Furthermore, studies on the interaction between ER and NF*κ*B in breast cancer cells suggest that ER represses NF*κ*B activity, suggesting a potential mechanism for estrogen’s anti-inflammatory activity in pre-menopausal women [56, 57]. The ER/NfKB interaction, however, is bidirectional and context-dependent [58]. Interestingly, both estrogen and androgen signalling show statistically significant change in correlation between Good and Bad clock groups.

**Figure 5:**
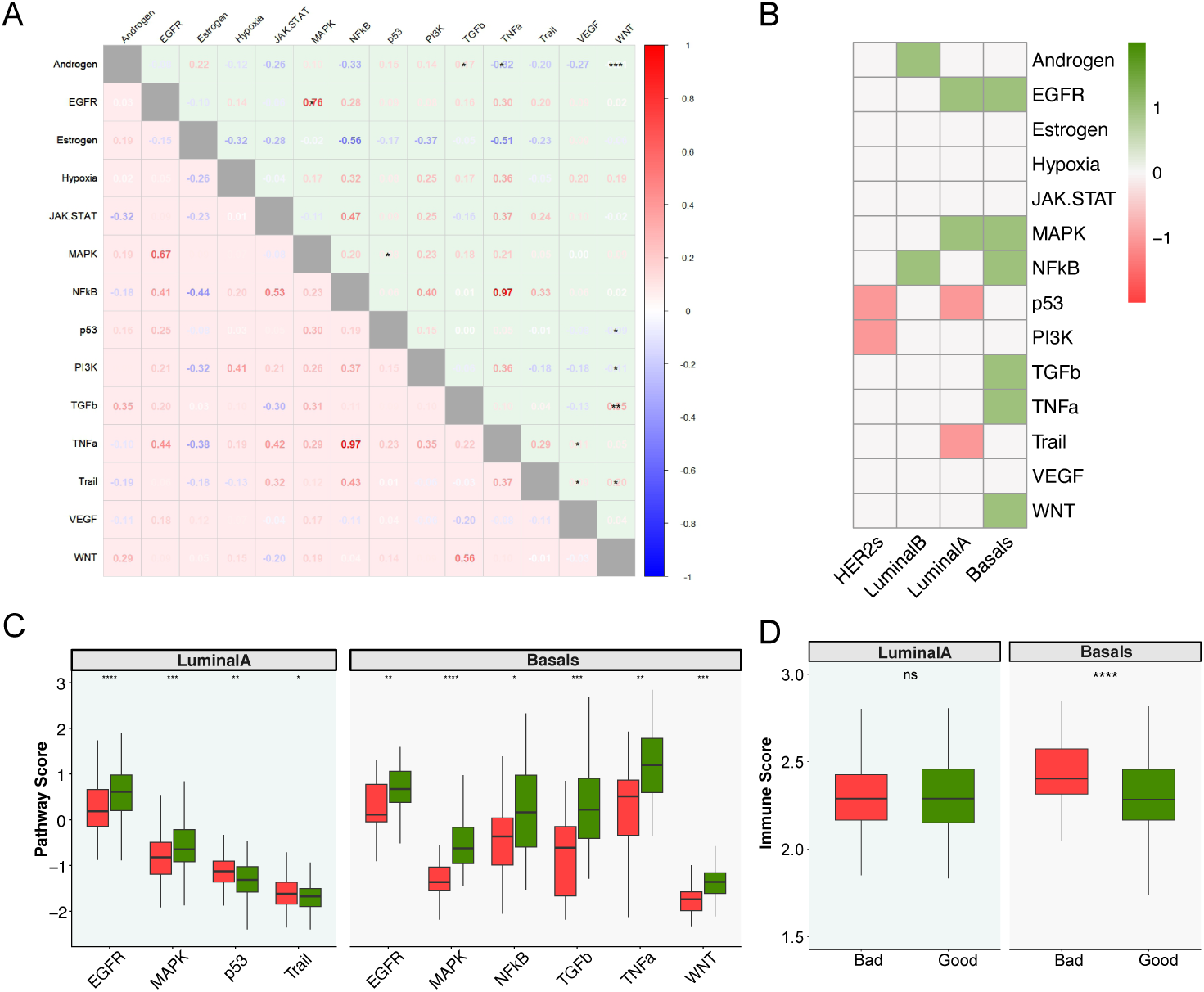
Θ-based stratification reveals distinct transcriptomic and pathway signatures between Good Clock Group (GCG) and Bad Clock Group (BCG). **A**. Correlation structure of PROGENy pathway activity scores between GCG (upper triangle) and BCG (lower triangle). Asterisks denote significant changes in pathway correlation structure between groups. Notable shifts in EGFR/MAPK and estrogen/androgen signalling correlations. **B**. Differential PROGENy pathway activation across PAM50 subtypes stratified by clock function. Green indicates pathways significantly more active in GCG; red indicates pathways significantly more active in BCG; gray indicates no significant difference. Basal-like tumours show most pronounced differences in important cancer-related pathways such as MAPK and TGF*β* signalling. **C**. Selected PROGENy pathway activity scores in Luminal A (least aggressive) and Basal-like (most aggressive) subtypes, stratified by clock function. GCG tumours show elevated pro-tumourigenic pathway activity compared to BCG, most pronounced in Basal-like subtype where EGFR, MAPK, TGF*β*, TNF*α*, and WNT are all significantly elevated. This pattern supports the counterintuitive finding that functional tumour clocks associate with worse survival by enabling coordinated activation of proliferative and metastatic signalling. **D**. Tumour immunogenicity assessed by Immunophenoscore in Luminal A and Basal-like subtypes. BCG Basal tumours show significantly higher immunogenicity compared to GCG, while no difference observed in Luminal A, suggesting clock dysfunction enhances immune recognition in aggressive subtypes. *Statistical note:* All pathway comparisons (panels E–I) performed using Wilcoxon rank-sum test. Significance levels indicated by asterisks: \**p <* 0.05, \*\**p <* 0.01, \*\*\**p <* 0.001, \*\*\*\**p <* 0.0001; ns = not significant.

We then used this method to examine changes in PROGENy pathways for all PAM50 subtypes in more detail because we have found in the survival analysis that there is significant difference between Luminal A and Basal subtypes’ response to Θ stratification. As expected, Figs. 5B-C and Fig. S19 suggest that the significantly differentially regulated pathways between Bad and Good Clock groups were PAM50 subtype-specific. Interestingly, Basals seem to exhibit the most pronounced differences in pathway activation (Fig. 5C). For example, MAPK is among the pathways with significantly reduced activation in the BCG. It is known that a dysfunctional circadian clock can lead to reduced activation of the MAPK pathway by disrupting the rhythmic expression of upstream regulators, such as growth factors and their receptors, which are tightly controlled by core clock components *BMAL1* and *CLOCK* [59, 60]. This loss of temporal regulation can result in attenuated or mistimed MAPK signalling. In the context of breast cancer, reduced MAPK pathway activation has been associated with less aggressive tumour behaviour and improved patient outcomes in certain subtypes [61–63]. Therefore, circadian disruption (even though generally considered detrimental) may, in specific molecular contexts, contribute to improved survival by dampening proliferative and pro-survival MAPK signalling in tumours.

While circadian disruption may protect against aggressive disease by reducing MAPK signalling, a contrasting mechanism emerges where functional clocks appear to facilitate worse outcomes through enhanced TGF*β* pathway coordination. It is known that TGF*β* signalling serves as an intercellular coupling factor that synchronises peripheral circadian oscillators through transcriptional regulation of core clock genes, and reciprocally, the core clock gene *BMAL1* is required for TGF*β*-induced pro-EMT signalling [64, 65]. In triple-negative and basal-like breast cancers, TGF*β* signalling is a well-established driver of epithelial-mesenchymal transition (EMT), tumour-initiating cell enrichment, and chemoresistance [66–68]. Importantly, EMT and cancer stem cell generation in breast cancer cells are temporally gated by intrinsic circadian clocks [69], suggesting that functional clocks enable precise temporal coordination of these aggressive phenotypes. The combination of functional circadian clocks with elevated TGF*β* signalling may create a particularly aggressive, self-reinforcing phenotype. TGF*β* signalling maintains clock function, while the intact clock reciprocally enables temporal coordination of TGF*β*-driven EMT and stress responses, supporting tumour cell survival under hostile microenvironmental conditions and enhancing both invasiveness and therapeutic resistance.

Another well studied feature of breast tumours shown to impact on patient survival is the tumour immune microenvironment [70, 71]. We investigated possible changes in the microenvironment using Immunophenoscore ( [72], SI Note S12), a computational algorithm developed to predict patient response to immune checkpoint inhibitor therapy that quantifies the immunogenicity of tumour samples by evaluating the expression of key genes indicative of effector cells such as CD8+ T cells and NK cells and immunosuppressive cells such as regulatory T cells and myeloid-derived suppressor cells, major histocompatibility complex molecules and immunomodulators (checkpoints and cytokines). Interestingly, we found a difference in immunogenicity between the TimeTeller GCG and BCG that was especially pronounced in PAM50 defined Basal tumours (Fig. 5D). This suggests that clock disruption in tumours is associated with higher immunogenicity in Basal tumours, opening an exciting possibility for stratifying effectiveness in immune checkpoint inhibitor drugs in the future. Moreover, this is in line with published results suggesting circadian disruption in peripheral tissues is associated with inflammation. Of note, no such difference can be detected in Luminal A tumours, consistent with the lack of a difference in BCG and GCG survival in this subtype.

Above we considered differences in downstream processes between tumours with Good and Bad Clocks. While identifying many potentially relevant pathways, further validation is needed to develop and test hypotheses on the precise nature of deregulation in Bad Clocks. We highlight the new possibilities that TimeTeller provides to study disease at the individual level across the spectrum of circadian dysfunction.

These findings provide further biological validation of Θ as a biomarker. Strong enrichment for circadian clock genes among DEGs independently confirmed that Θ measures circadian clock function, not an unrelated process. The large number of DEGs (≈1,200 genes) and pathway-level changes demonstrated that Θ reflects system-wide alterations in tumour biology, not merely expression changes in the training genes used to calculate it. These extensive transcriptomic differences between high- and low-Θ tumours help explain Θ’s prognostic value for survival and suggest that chronotherapy or direct circadian targeting might affect the two groups differently. Critically, Θ stratification identifies biologically meaningful tumour subtypes undetected by conventional molecular classification, providing orthogonal information for patient stratification and therapeutic targeting.

In summary, the Θ stratification identifies biologically meaningful tumour subtypes that would not be detected by conventional molecular classification schemes. This provided orthogonal information for patient stratification and potential therapeutic targeting.

### Conclusion and discussion

This study further establishes TimeTeller as a novel single-sample circadian biomarker that quantifies both internal circadian phase and clock dysfunction in individual patient tumours. Applied to 1,286 breast cancer patients across six clinical cohorts, our analysis revealed several key findings that challenge conventional understanding of circadian biology in cancer. By demonstrating its utility in a major human health context, it greatly extends the results in [38], where it was validated against diverse murine datasets.

Our first major finding is that most breast tumours retain mechanistically functional circadian clocks. This is evidenced by the near-linear relationship between TT time and the multidimensional clock gene expression state, coherent phase predictions, and appropriate gene-gene correlations across the circadian network. However, these clocks display critical alterations: they are temporally uncoupled from external time of day, and both tumour internal circadian phases and the corresponding clock gene states are compressed compared to what would be expected in a healthy clock.

Our most striking finding challenges the assumption that circadian disruption uniformly promotes cancer progression. For the approximately 70-80% of all patients with the lowest dysfunction score, increasing circadian dysfunction (higher dysfunction score) associates with improved survival across multiple endpoints and study centers. This effect is particularly pronounced in large tumours (T3/4), where relative risk decreases nearly five-fold, and in aggressive molecular subtypes (Basal and Luminal B). The prognostic value of Θ is independent of treatment status and proliferation markers.

These results are independent of treatment, suggesting that both the altered clock and Θ reflect intrinsic tumour biology rather than treatment response and could be used to stratify patients in both treated and untreated populations.

As mentioned above, this finding has partial precedent: Li and colleagues recently reported similar results for Luminal A breast cancer patients using CYCLOPS2.0 in TCGA data [26]. Given their fundamentally different methods (CYCLOPS2.0 algorithm using low amplitude at the population level as corresponding to dysfunction) and different data (TCGA), it is notable that they also found non-monotonic dependence of survival on dysfunction level, suggesting this represents a fundamental biological phenomenon. However, their analysis allocated dysfunction scores to groups (lowest, intermediate, and highest amplitude) rather than to individuals.

These findings support a provocative reinterpretation of the nature of the impact of disruption on the survival of patients. Rather than representing simple collateral damage from oncogenesis, they suggest that in the majority of breast cancer patients, the tumour circadian clock has been actively reconfigured into a mechanistically functional oscillator that supports cancer progression and thereby reduces patient survival. This concept reframes circadian disruption as an active, perhaps selected trait of tumour evolution rather than merely a consequence of malignant transformation. Moreover, the paradoxical protective effect of dysfunction suggested therapeutic strategies should disrupt rather than restore tumour circadian function—a significant conceptual shift with implications for chronotherapy and treatment design.

In fact, targetting the clock pharmacologically to treat cancer has previously been proposed [73, 74], and at least one such small molecule, SHP1705, has already been reported to be in clinical phase I studies [75].

Our results align with and extend the emerging concept of circadian clock hijacking in cancer [26, 76–78]. Recent work has shown that tumour cells can exploit circadian rhythms to enhance their survival and dissemination through multiple mechanisms. Cancer cells may time their immune invisibility in coordination with host circadian rhythms [76–78], usurp circadian transcriptional machinery to reprogram metabolic processes for optimal proliferation conditions [79–81], and suppress apoptosis or schedule therapeutic resistance to coincide with specific circadian phases [26, 82]. The metastatic cascade itself may be temporally organized and linked to circadian machinery [26, 83].

The independence of Θ’s prognostic value from conventional markers makes it especially valuable for risk stratification in populations where existing tools perform poorly, particularly in Basal (predominantly triple-negative) and Luminal B subtypes. TimeTeller offers several practical advantages for clinical implementation. As a single-sample method, it does not require population reference data or paired samples, making individualized assessment feasible in routine clinical settings. Beyond prognostics, Θ stratification reveals an underappreciated axis of tumour biology orthogonal to conventional molecular subtyping, providing a framework for mechanistic investigation of how circadian state interacts with oncogenic drivers, immune infiltration, and therapeutic vulnerabilities.

The current TimeTeller implementation requires training on circadian time-series data, which for human data currently limits us to microarray platforms because comprehensive healthy human RNA-seq circadian time-series datasets are not yet available. Although TimeTeller has been successfully applied to murine and primate RNA-seq data, until such human RNA datasets exist this restricts direct application to modern RNA-seq based clinical assays, though we are actively developing approaches for RNA-seq platforms and exploring training-free implementations that could circumvent this limitation. Moreover, some limitations warrant careful consideration.

#### Tumour heterogeneity

Bulk tissue analysis complicates interpretation of our results. It remains fundamentally unclear whether Θ primarily reflects cancer cell properties, contributions from mixed cell populations within the tumour, or microenvironmental factors such as stromal or immune cell composition. Single-cell and spatial transcriptomics have unveiled a significant impact of the tumour microenvironment on tumour biology and treatment outcomes [84, 85]. These approaches could clarify cell-type specific contributions to the Θ signal and determine whether circadian dysfunction is uniform across the tumour or exhibits spatial heterogeneity.

#### Retrospective nature and validation

While our analysis spans six clinical cohorts with consistent results, it remains fundamentally retrospective. Prospective validation in independent patient cohorts is essential to confirm clinical utility and establish Θ thresholds for clinical decision-making. Prospective studies would also enable assessment of whether Θ measurements are stable over time or change during disease progression and treatment. Importantly, such studies could also collect data about the patient’s overall rhythm, as it has been shown that breast cancer can disrupt cortisol rhythms in patients, which might have implications for the CD8+ T cell invasion of the tumour as recently suggested using a mouse model [86]

#### Mechanistic understanding

The mechanistic basis for the protective effect of circadian dysfunction remains incompletely understood and requires functional validation. While we propose several plausible mechanisms (metabolic flexibility, immune evasion, metastatic timing), experimental studies in cell culture and animal models are needed to establish causal relationships and identify specific molecular pathways linking clock dysfunction to reduced tumour aggressiveness. Understanding these mechanisms is critical for rational design of clock-disrupting therapeutic interventions.

#### Generalizability

Our study focused exclusively on breast cancer, and generalizability to other tumour types remains unknown. Different cancer types may exhibit distinct relationships between circadian function and progression depending on their tissue of origin, molecular drivers, and microenvironmental characteristics. Systematic evaluation across tumour types is needed to determine whether clock reconfiguration is a pan-cancer phenomenon or specific to breast cancer.

#### Nonlinearity and subgroup effects

The nonlinear relationship between Θ and survival, with potential risk increase at very high Θ values (though not statistically significant), warrants further investigation in larger cohorts. The limited number of patients with very high Θ values (*>* 0.4) constrains our ability to definitively characterize this region. Understanding whether severely disrupted clocks behave differently from moderately disrupted clocks has important implications for therapeutic strategies.

In summary, as precision oncology advances toward increasingly personalized treatment selection, incorporating temporal dimensions becomes essential. The fundamental question raised by this work extends beyond breast cancer: should treatment strategies now consider temporal disruption as a therapeutic goal? If tumours with disrupted circadian organization are less aggressive, developing therapies that disrupt tumour circadian machinery while preserving host rhythms may represent an unexplored opportunity. TimeTeller provides the measurement framework to test this hypothesis and guide such interventions toward clinical reality.

## Ethics statement

The study associated with the Bjarnason et al. human training data [87] was approved by the Sunnybrook Health Sciences Centre Research Ethics Board. Project identification number 396–2004. Written informed consent was obtained from each subject as requested by the Research Ethics Board.

## Author Contributions

**Conceptualisation and project initialisation**: Sylvie Giacchetti, Francis Levi, David A. Rand, Denise Vlachou.

**Methodology**: David A. Rand, Vadim Vasilyev, Denise Vlachou.

**Data curation**: Georg A. Bjarnason, Vadim Vasilyev.

**Investigation**: Georg A. Bjarnason, Sylvie Giacchetti, Tami A. Martino.

**Formal analysis**: Vadim Vasilyev, Denise Vlachou.

**Funding acquisition**: Robert Dallmann, David A. Rand.

**Supervision**: Robert Dallmann, David A. Rand.

**Writing – original draft & editing**: Robert Dallmann, Vadim Vasilyev, David A. Rand.

**Writing – review & editing**: Francis Levi, Sylvie Giacchetti.

