## Supplementary Information for "A robust negative association between estimated tumour circadian clock function and survival in early stage breast cancer"

### Supplementary Material for the paper "A robust negative association between estimated tumour circadian clock function and survival in early stage breast cancer"

Vadim Vasilyev[1]* Denise Vlachou[2]† Sylvie Giacchetti[3]‡
Georg A. Bjarnason[4]§ Tami A. Martino[5]¶ Francis Levi[6]‖ Robert Dallmann[1]**
David A Rand[7]††

[1]Division of Biomedical Sciences, Warwick Medical School, University of Warwick, Coventry, UK
[2]GSK Research, Gunnels Wood Road, Stevenage, Hertfordshire, UK
[3] APHP, Senology Unit, Hospital Saint Louis, Paris, France; INSERM Unit 1193 HEPAREG, and Faculty of Medicine, Paris Saclay University, France
[4]Odette Cancer Centre, Sunnybrook Health Sciences Centre, Faculty of Medicine. University of Toronto. Toronto, ON, M4N 3M5, Canada
[5] Biomedical Sciences, University of Guelph, Guelph, Ontario, Canada, N1G 2W1
[6]INSERM Unit 1193 HEPAREG, and Faculty of Medicine, Paris-Saclay University, Paul Brousse hospital, Villejuif, France Gastro-intestinal and Medical Oncology Department, Paul-Brousse Hospital, Assistance Publique - Hôpitaux de Paris, Villejuif, France Department of Statistics, University of Warwick, Coventry, UK
[7]Mathematics Institute & Zeeman Institute for Systems Biology and Infectious Disease Epidemiology Research, University of Warwick, Coventry, UK

#### Contents

**I Supplementary Notes** S3

**S1 Human datasets used** S3

**S2 Microarray data pre-processing** S4

**S3 Data Processing and Analysis** S4

**S4 Normalisation and LogThresh selection** S5

**S5 Day/Night Classification Using Lasso Logistic Regression** S5

**S6 Population-level evidence confirms oscillatory rather than arrhythmic clocks** S5

*
†
‡
§
¶
‖

**S7 PAM50 classification** **S6**

**S8 Survival Analysis** **S6**

**S9 Proliferation Index** **S7**

**S10 Differential Expression** **S8**

**S11 Pathway Analysis** **S8**

**S12 Immunophenotyping** **S8**

**II Supplementary Tables** **S8**

**III Supplementary Figures** **S17**

### Part I
### Supplementary Notes

Throughout these notes the main paper is referred to as **I**.

#### S1 Human datasets used

| **Human training dataset** | |
|---|---|
| **Bjarnason *et al.*** (*1*) | |
| Technology | Affymetrix Human Genome U133 Plus 2.0 array |
| GEO | |
| Tissue(s) | Oral mucosa |
| Samples | 10 healthy volunteers |
| Study | Investigation of diurnal rhythms in gene expression in human oral mucosa collected from healthy subjects and potential implications for gender differences in toxicity, response and survival and optimal timing of targeted therapy |
| Timepoints | 6 time points (every 4 hours) |
| Number of replicates | 1 |
| Gender | 5 male and 5 female |

| **Breast cancer datasets** | |
|---|---|
| **REMAGUS02** (*2*) | |
| Technology | Microarray (Affymetrix Human Genome U133 Plus 2.0 Array) |
| GEO | GSE26639 |
| Tissue(s) | Breast tumors biopsies |
| OS available | Yes |
| DMFS available | No |
| RFS available | Yes |
| Number of patients | 226 |
| Treatment | Yes (226) |
| **Upp** (*3*) | |
| Technology | Microarray (Affymetrix Human Genome U133A, U133B Array) |
| GEO | GSE3494 |
| Tissue(s) | Breast tumors biopsies |
| OS available | No |
| DMFS available | No |
| RFS available | Yes |
| Number of patients | 233 |
| Treatment | No (157), Yes (76) |
| **Unt** (*4*) | |
| Technology | Microarray (Affymetrix Human Genome U133A Array) |
| GEO | GSE2990 |
| Tissue(s) | Breast tumors biopsies |
| OS available | No |
| DMFS available | Yes |
| RFS available | Yes |
| Number of patients | 125 |
| Treatment | No (125) |
| **Mainz** (*5*) | |
| Technology | Microarray (Affymetrix Human Genome U133A Array) |
| GEO | GSE11121 |
| Tissue(s) | Breast tumors biopsies |
| OS available | No |
| DMFS available | Yes |
| RFS available | No |
| Number of patients | 200 |

| Breast cancer datasets (cont'd) | |
|---|---|
| Treatment | No (200) |
| **Rotterdam** (*6*) | |
| Technology | Microarray (Affymetrix Human Genome U133A Array) |
| GEO | GSE2034 |
| Tissue(s) | Breast tumors biopsies |
| OS available | No |
| DMFS available | Yes |
| RFS available | No |
| Number of patients | 286 |
| Treatment | No (286) |
| **Transbig** (*7*) | |
| Technology | Microarray (Affymetrix Human Genome U133A Array) |
| GEO | GSE7390 |
| Tissue(s) | Breast tumors biopsies |
| OS available | Yes |
| DMFS available | Yes |
| RFS available | Yes |
| Number of patients | 198 |
| Treatment | No (198) |

#### S2 Microarray data pre-processing

For Affymetrix GeneChip data, raw CEL files were obtained, and the *AffyBatch* object (*affy* package v. 1.82.0 of Bioconductor v. 3.19 in R 4.4.1) was preprocessed using the fRMA method (*8*) to minimise batch effects when analysing multiple studies. Probes were filtered for multiple known gene matches and the resulting *ExpressionSet* object was reduced to the set of common probes (Affymetrix Human Genome U133A Array being a subset of Affymetrix Human Genome U133 Plus 2.0 Array). GSM accession numbers were checked for all the samples and the duplicates were removed from the *Unt* cohort that also appeared in the *Upp* cohort.

Where necessary for downstream analysis, multiple microarray probes for a single gene were collapsed using the *MaxMean* method of the *collapseRows* function from *WGCNA* package (v. 1.73) and then merged by the gene identifier. This was done due to the evidence that choosing the probe with the highest average expression across samples leads to best between-study consistency (*9*).

#### S3 Data Processing and Analysis

The Bioconductor packages *breastCancerUNT*, *breastCancerUPP*, *breastCancerTRANSBIG*, and *breastCancerMAINZ* were used to obtain clinical information. Where available, numerical tumor sizes were classified as: T1 (0–2 cm], T2 (2–5 cm], T3 (5–15 cm]. Time to event was converted to days, and right censoring was applied at 10 years for patients without events.

Statistical analyses used the *survival* v. 3.7-0 and *rms* v. 6.8-2 R packages. Survival curves were constructed using the Kaplan-Meier method, and significance was assessed using the log-rank test. Univariate and multivariate prognostic factors were analyzed using Cox proportional hazards models (*10*). The proportional hazards assumption was assessed by visual examination of log(−log) survival plots and testing scaled Schoenfeld residuals using the *cox.zph* function. After stratifying by PAM50 subtype, the assumption held for all models.

Statistical significance (global effects, fixed effects, and interactions) was assessed using likelihood ratio tests. Models were corrected for optimism using internal validation with 300 bootstrap repetitions via the *rms* package. Reported Somers' $D_{xy}$ values (and the linearly related C-index) are therefore optimism-corrected and unbiased.

#### S4 Normalisation and LogThresh selection

Intergene normalisation is used in this work, since test data comes in a format of single sample per patient. For a REV $g = (g_i)$, this produces normalised levels $\hat{g}_i = (g_i - \mu)/\sigma$, where $\mu$ and $\sigma^2$ are the mean and variance of the $g_i$ values. This effectively maps each REV onto its shape as a unit vector.

The choice of $l_{\text{thresh}}$ should be as large as possible while satisfying: (i) very few training and control samples have flat regions significantly intersecting $C(t|T)$ that contribute to $\Theta$, and (ii) most test samples have MLs above $exp(l_{\text{thresh}})$.

Two considerations prevent decreasing $l_{\text{thresh}}$ further. First, we must protect against inaccurate exceptionally low $L_{g,i}$ values. Second, excessive reduction removes meaningful LRF structure away from time $T$. This is because decreasing $l_{\text{thresh}}$ reduces $L_g(t)$ more at times far from $T$ than near $T$, flattening the LRF away from the peak. Whether dysfunction manifests as low ML or secondary peaks, over-reduction of $l_{\text{thresh}}$ moves these features below $C(t|T)$, decreasing $\Theta$.

When $\log \text{ML}$ is marginally above $l_{\text{thresh}}$, the resulting flat intervals above $C(t|T)$ contribute to $\Theta$, providing information about how low the maximum ML is.

If criterion (ii) removes excessive structure, $l_{\text{thresh}}$ may be increased provided criterion (i) violations remain limited. Similarly, too many test samples violating (ii) is undesirable, as these have $\Theta = 1$, lacking non-trivial stratification despite indicating higher dysfunction.

#### S5 Day/Night Classification Using Lasso Logistic Regression

To assess the ability of clock gene expression patterns to discriminate between day and night phases, we employed a regularized logistic regression approach. All unique pairwise ratios (n = 45) were computed from the filtered rhythmic clock probes (n = 10 probes), generating a feature set of gene expression ratios for each sample. Pairwise ratios were used to capture relative expression relationships between clock genes while normalizing for sample-specific technical variation.

Samples were assigned binary labels based on collection time, with times between 21:00 and 07:00 classified as ”night” and all other times as ”day”. Lasso (Least Absolute Shrinkage and Selection Operator) logistic regression was performed using the *glmnet* package (v.4.1) in *R* to identify the minimal set of gene expression ratios capable of discriminating day from night phases. The regularization parameter $\lambda$ was optimized using 10-fold cross-validation, selecting the value that minimized classification error (*lambda.min*).

The final model selected 5 pairwise ratio features and achieved 97% classification accuracy on the training data, indicating that a sparse subset of clock gene relationships is sufficient to capture day/night phase information.

#### S6 Population-level evidence confirms oscillatory rather than arrhythmic clocks

In addition to the single sample analysis given in the main paper **I** verifing that circadian clocks remain functionally oscillating diurnally in our tumor samples, we also tested this using population methods similar to those in (*11*). This previous work on untimed single samples confirmed clock oscillation by examining gene-gene correlations among core clock genes and fitting nonlinear regression to clock gene expression versus estimated time. While not definitively ruling out clock arrest, such coherent population-level patterns are incompatible with randomly frozen or arrhythmic clocks.

We performed three complementary analyses to assess clock oscillation in our tumor data. First, we compared gene-gene correlations for core clock genes between training data and tumor samples stratified by quartile (Fig. S17a,b). Functional oscillators should show positive correlations between transcriptional activators and negative correlations with repressors. Second, we plotted expression levels of the 9 clock probes against their estimated TT phase for each sample (SI Figs. S4 & S5), revealing coherent time dependence compatible with the behaviour of the probe in the training clock.

Third, we verified that the observed nREVs and associated TT times of tumor samples reflected those of the training clock, with accuracy improving as model confidence (ML) increased (SI Fig. S8b).

All three analyses demonstrate population-level gene-gene correlations matching patterns expected from functional oscillators. Combined with single-sample evidence that TT times fall predominantly during working hours and that gene expression varies coherently with estimated phase, these findings strongly support maintained oscillatory clock function in tumors.

### S7 PAM50 classification

The PAM50 classifier for identification of breast cancer molecular subtypes was used to improve the prognostic model during the multivariate survival analysis (*12*). For that *genefu* package v. 2.36.0 was used with the *pam50.robust* model chosen (official centroids with robust scaling of the gene expressions). This was shown to achieve the best concordance with the standard clinically used biomarkers such as ER immunohistochemistry, HER2 IHC fluorescence in situ hybridization (FISH) and histological grade. The model was chosen, since in the combined cohort we do not expect any sampling bias with all the 5 subtypes present.

| | Basals | HER2s | LuminalA | LuminalB | Normals |
|---|---|---|---|---|---|
| ER$^-$PGR$^-$HER2$^-$ | 37 | 2 | 3 | 4 | 2 |
| ER$^-$PGR$^-$HER2$^+$ | 1 | 24 | 3 | 4 | 0 |
| ER$^-$PGR$^+$HER2$^-$ | 0 | 0 | 1 | 0 | 0 |
| ER$^-$PGR$^+$HER2$^+$ | 0 | 1 | 1 | 1 | 0 |
| ER$^+$PGR$^-$HER2$^-$ | 2 | 0 | 8 | 17 | 0 |
| ER$^+$PGR$^-$HER2$^+$ | 0 | 4 | 0 | 11 | 0 |
| ER$^+$PGR$^+$HER2$^-$ | 0 | 0 | 33 | 28 | 0 |
| ER$^+$PGR$^+$HER2$^+$ | 0 | 6 | 11 | 12 | 0 |

**Figure S1:** Comparing PAM50 categories against IHC classification for the REMAGUS patients.

The PAM50 gene expression classifier stratifies breast cancers into five intrinsic molecular subtypes based on a 50-gene signature. These subtypes include: Luminal A (ER-associated gene expression); Luminal B (similar to Luminal A but with higher proliferative gene expression); HER2-enriched (elevated ERBB2 and HER2 pathway genes); Basal-like (predominantly triple-negative with high basal cytokeratin, EGFR, and proliferation marker expression); and Normal-like (resembling normal breast tissue). The PAM50 classification demonstrated strong concordance with clinical biomarkers including immunohistochemistry-based oestrogen receptor and HER2 status, and histological grade, providing robust prognostic stratification in breast cancer.

### S8 Survival Analysis

Linear modeling, while superior to dichotomization, assumes constant marginal effects regardless of predictor magnitude (*13, 14*). However, many biological relationships exhibit threshold effects, plateaus, or other non-linearities that linear models cannot capture (**?**, *14*).

We employed a powerful Cox proportional hazards model (*13*) that treats $\Theta$ as a continuous variable while adjusting for appropriate covariates. In the standard Cox model, the log hazard function is $\log \lambda(t|X) = \beta_1 \Theta + \sum_{i>1} \beta_i X_i$, which is linear in $\Theta$ and other covariates $X_i$. When this linearity assumption is questionable for continuous covariates such as $\Theta$, restricted cubic spline functions allow detection and characterization of nonlinear effects (*13*). In our analysis, this approach uncovered important patterns that linear analysis missed.

In the nonlinear Cox model, we replace the linear term $\beta_1 \Theta$ with a nonlinear spline function $f(\Theta)$, enabling formal testing of whether the nonlinearity significantly improves model fit (see below).

##### Restricted Cubic Splines

For a continuous predictor $X$ with $k$ knots at $t_1, \ldots, t_k$, the restricted cubic spline function is (*13, 14*):

$$f(X) = \beta_0 + \beta_1 X + \sum_{j=2}^{k-1} \beta_j X_j \tag{1}$$

where the nonlinear basis functions ($j = 2, \ldots, k-1$) are:

$$X_j = (X - t_{j-1})_+^3 - (X - t_{k-1})_+^3 \frac{(t_k - t_{j-1})}{(t_k - t_{k-1})} + (X - t_k)_+^3 \frac{(t_{k-1} - t_{j-1})}{(t_k - t_{k-1})} \tag{2}$$

with $(u)_+ = \max(0, u)$. This formulation ensures smooth joins (continuous first and second derivatives) at knots while constraining $f(X)$ to be linear in the tails ($X \leq t_1$ and $X \geq t_k$) (*13, 14*).

With $k$ knots, only $k - 1$ parameters (plus intercept) require estimation, compared to $k + 3$ for unrestricted cubic splines (*13*). Nonlinearity can be tested via a likelihood ratio test of the null hypothesis $H_0 : \beta_2 = \ldots = \beta_{k-1} = 0$. If significant, the nonlinear terms are retained in the model. The linear tail constraint prevents unexpected behavior in sparse data regions, which is particularly valuable for skewed distributions (*14, 15*).

**Key advantages**: (i) data-driven functional form discovery without arbitrary constraints (*13–15*); (ii) efficient parameter usage—a three-knot spline uses the same degrees of freedom as quartile categorization while capturing smooth transitions (*14*); (iii) standard inferential methods remain valid (*13–15*); (iv) predictions available at any predictor value (*13, 14*).

##### Implementation

We employed three-knot restricted cubic splines ($k = 3$) with knots at the 5th, 50th, and 95th percentiles (similar results were obtained with $k = 4$, saving one degree of freedom). This specification requires two degrees of freedom (one linear, one nonlinear basis function) and adequately captures common nonlinear patterns while conserving parameters (*14, 15*). Quantile-based knot placement ensures adequate sample sizes per region and robustness to outliers (*13*). Basis functions were normalized by $\tau = (t_3 - t_1)^2$ for numerical stability (*13*).

Since the $\Theta$ distribution often exhibits right skewness, restricted cubic splines were particularly appropriate due to their linear tail constraints, which improve performance in regions with sparse data.

#### S9 Proliferation Index

Single-sample Gene Set Enrichment Analysis (ssGSEA) was used to quantify pathway activity independently for each sample (*16*). Unlike standard GSEA, ssGSEA calculates enrichment scores for each sample-gene set pair without requiring phenotype labels, estimating the degree to which genes in a pathway are coordinately up- or down-regulated within individual samples.

**Table S3:** Core proliferation marker genes (n=25) used for calculating pseudo-proliferation index.

| Functional Group | Genes |
|---|---|
| MCM Complex | *MCM2*, *MCM3*, *MCM4*, *MCM5*, *MCM6*, *MCM7* |
| Proliferation Markers | *PCNA*, *MKI67*, *PLK1* |
| DNA Replication | *FEN1*, *RRM1*, *RRM2*, *CDK1*, *PRIM2*, *POLA1* |
| RFC Complex | *RFC2*, *RFC3*, *RFC4*, *RFC5* |
| RPA Complex | *RPA1*, *RPA2*, *RPA3* |
| Cohesin Complex | *SMC1A*, *SMC3*, *STAG2* |

To assess tumor proliferative capacity, we used a 25-gene pseudo-proliferation index derived by Locard-Paulet et al. (*17*) from analysis of NCI60 cell lines with known doubling times. This index comprises established proliferation markers (*MCM2-7* complex, *CDK1*, *PCNA*, *PLK1*, *MKI67*), genes involved in DNA replication machinery (*RPA1-3*, *RFC2-5*, *RRM1-2*, *FEN1*, *PRIM2*, *POLA1*), and chromosome segregation (*SMC1A*, *SMC3*, *STAG2*), selected for their consistent correlation with cell doubling times across multiple proteomics and transcriptomics datasets.

ssGSEA was performed using the *hacksig* package v.0.1.2 (*18*). Enrichment scores were normalized to $[0, 1]$ independently across samples, and the weighting exponent $\alpha$ was set to 0.25 (default).

#### S10 Differential Expression

Differential expression analysis was performed using the *limma* package v.3.60.4 (*19*). Principal component analysis of the 3,000 most variable features was used to identify relevant covariates. The linear model included circadian dysfunction (categorical split into Good, Average and Bad), ER status, PAM50 subtype, and dataset (study) as covariates. Batch effects were addressed by including dataset as a covariate in the linear model rather than applying ComBat batch correction, following *limma* user guide recommendations. To account for variable RNA sample quality across patient samples, sample-specific weights were estimated using *limma*'s *arrayWeights* function. Differentially expressed genes were identified following empirical Bayes moderation and Benjamini-Hochberg multiple testing correction (adjusted $p < 0.05$).

To assess statistical robustness, permutation analysis was performed using the same model structure but with randomly permuted circadian dysfunction group labels.

The REMAGUS02 dataset was analyzed separately due to substantial differences in count distributions between microarray platforms (U133 Plus 2.0 vs U133A) that could not be adequately addressed through covariate adjustment.

#### S11 Pathway Analysis

Pathway analysis was conducted using two complementary approaches. Standard enrichment analysis was performed using the *gprofiler2* package v.0.2.3 (*20*) with *ordered_query* = *TRUE*, supplying differentially regulated genes ranked by $\log_2$ fold change.

To complement gene set enrichment with pathway activity inference, we used the *progeny* package v.1.26.0 (*21*). *Progeny* estimates pathway activity from gene expression by leveraging a compendium of perturbation experiments to identify Pathway RespOnsive GENes (PROGENy) that are consistently deregulated across multiple stimuli. This approach distinguishes between pathway-associated gene expression and functional pathway activation. The model matrix was constructed using the top 100 most responsive genes (default), and statistical significance was assessed using 200 permutations with a gene sampling-based permutation strategy.

#### S12 Immunophenotyping

Tumor immunogenicity was quantified using the Immunophenoscore (IPS), calculated using the *hacksig* R package (v0.1.2) (`hack_immunophenoscore()` function). The IPS integrates expression of genes representing four determinants of tumor immunogenicity identified through machine learning analysis of TCGA pan-cancer data (*22*): (i) MHC molecules (antigen processing machinery), (ii) immunomodulators (checkpoint and co-stimulatory molecules), (iii) effector cells (activated and effector memory $CD4^+$/$CD8^+$ T cells), and (iv) suppressor cells (regulatory T cells and myeloid-derived suppressor cells). Raw IPS values, representing the sum of weighted Z-scores across categories (positive weights for immunostimulatory factors, negative for immunosuppressive), were used for statistical comparisons.

### Part II
### Supplementary Tables

**Table S1:** Clinical characteristics and endpoint data availability by cohort

| Cohort | No. Patients | OS | RFS | DMFS | Treatment | T3/4[a] |
|---|---|---|---|---|---|---|
| **Remagus** | 226 | ✓ | ✓ | ✗ | Yes | 105 |
| **TRANSBIG** | 198 | ✓ | ✓ | ✓ | No | 0 |
| **Mainz** | 200 | ✗ | ✗ | ✓ | No | 3 |
| **Upp** | 251 | ✗ | ✓ | ✗ | Mixed | 6 |
| **Unt** | 125 | ✗ | ✓ | ✓ | No | 1 |
| **Rot** | 286 | ✗ | ✗ | ✓ | No | NA |

[a] Number of T3 and T4 tumors combined.

**Table S2:** Overall Survival - Patient Characteristics

| Overall Survival | | | | | |
|---|---|---|---|---|---|
| | | | Clock Function | | |
| **Variable** | **N** | **Overall**, N = 411[1] | **Good**, N = 81[1] | **Average**, N = 276[1] | **Bad**, N = 54[1] |
| **Theta** | 411 | 0.27 (0.18) | 0.06 (0.02) | 0.27 (0.10) | 0.60 (0.12) |
| **Age** | 411 | 46.84 (8.14) | 46.53 (8.62) | 46.76 (8.08) | 47.72 (7.74) |
| **Treatment** | 411 | | | | |
| No Treatment | | 197 (48%) | 30 (37%) | 153 (55%) | 14 (26%) |
| Treatment | | 214 (52%) | 51 (63%) | 123 (45%) | 40 (74%) |
| **Tumour Size** | 411 | | | | |
| T1 | | 101 (25%) | 13 (16%) | 79 (29%) | 9 (17%) |
| T2 | | 209 (51%) | 38 (47%) | 146 (53%) | 25 (46%) |
| T3_4 | | 101 (25%) | 30 (37%) | 51 (18%) | 20 (37%) |
| **ER Status** | 411 | | | | |
| 0 | | 148 (36%) | 25 (31%) | 104 (38%) | 19 (35%) |
| 1 | | 263 (64%) | 56 (69%) | 172 (62%) | 35 (65%) |
| **Grade** | 402 | | | | |
| 1 | | 43 (11%) | 8 (11%) | 30 (11%) | 5 (9.3%) |
| 2 | | 162 (40%) | 36 (47%) | 104 (38%) | 22 (41%) |
| 3 | | 197 (49%) | 32 (42%) | 138 (51%) | 27 (50%) |
| (Missing) | | 9 | 5 | 4 | 0 |
| **PAM50** | 411 | | | | |
| Basals | | 84 (20%) | 13 (16%) | 65 (24%) | 6 (11%) |
| HER2s | | 57 (14%) | 10 (12%) | 39 (14%) | 8 (15%) |
| LuminalA | | 140 (34%) | 37 (46%) | 89 (32%) | 14 (26%) |
| LuminalB | | 119 (29%) | 16 (20%) | 78 (28%) | 25 (46%) |
| Normals | | 11 (2.7%) | 5 (6.2%) | 5 (1.8%) | 1 (1.9%) |
| **Centre** | 411 | | | | |
| Rem_Cent_1 | | 105 (26%) | 24 (30%) | 64 (23%) | 17 (31%) |
| Rem_Cent_2 | | 58 (14%) | 13 (16%) | 35 (13%) | 10 (19%) |
| Rem_Cent_3 | | 17 (4.1%) | 4 (4.9%) | 10 (3.6%) | 3 (5.6%) |
| Rem_Cent_4 | | 34 (8.3%) | 10 (12%) | 14 (5.1%) | 10 (19%) |
| TRANSBIG | | 197 (48%) | 30 (37%) | 153 (55%) | 14 (26%) |

[1] Mean (SD); n (%)

**Table S3:** Distant Metastasis Free Survival - Patient Characteristics

| Distant Metastasis Free Survival | | | | | |
|---|---|---|---|---|---|
| | | | **Clock Function** | | |
| **Variable** | **N** | **Overall**, N = 789[1] | **Good**, N = 141[1] | **Average**, N = 486[1] | **Bad**, N = 162[1] |
| **Theta** | 789 | 0.31 (0.22) | 0.06 (0.02) | 0.27 (0.10) | 0.65 (0.14) |
| **Age** | 512 | 52.92 (11.67) | 51.78 (9.95) | 52.03 (11.97) | 56.66 (11.43) |
| (Missing) | | 277 | 53 | 165 | 59 |
| **Tumour Size** | 511 | | | | |
| T1 | | 285 (56%) | 48 (55%) | 178 (55%) | 59 (57%) |
| T2 | | 222 (43%) | 39 (45%) | 140 (44%) | 43 (42%) |
| T3 | | 4 (0.8%) | 0 (0%) | 3 (0.9%) | 1 (1.0%) |
| (Missing) | | 278 | 54 | 165 | 59 |
| **ER Status** | 783 | | | | |
| 0 | | 210 (27%) | 49 (35%) | 140 (29%) | 21 (13%) |
| 1 | | 573 (73%) | 90 (65%) | 343 (71%) | 140 (87%) |
| (Missing) | | 6 | 2 | 3 | 1 |
| **Grade** | 494 | | | | |
| 1 | | 89 (18%) | 10 (13%) | 63 (20%) | 16 (16%) |
| 2 | | 262 (53%) | 38 (48%) | 155 (50%) | 69 (68%) |
| 3 | | 143 (29%) | 31 (39%) | 95 (30%) | 17 (17%) |
| (Missing) | | 295 | 62 | 173 | 60 |
| **PAM50** | 789 | | | | |
| Basals | | 137 (17%) | 36 (26%) | 89 (18%) | 12 (7.4%) |
| HER2s | | 81 (10%) | 12 (8.5%) | 55 (11%) | 14 (8.6%) |
| LuminalA | | 359 (46%) | 56 (40%) | 215 (44%) | 88 (54%) |
| LuminalB | | 174 (22%) | 24 (17%) | 105 (22%) | 45 (28%) |
| Normals | | 38 (4.8%) | 13 (9.2%) | 22 (4.5%) | 3 (1.9%) |
| **Centre** | 789 | | | | |
| MAINZ | | 191 (24%) | 11 (7.8%) | 100 (21%) | 80 (49%) |
| Rot | | 277 (35%) | 53 (38%) | 165 (34%) | 59 (36%) |
| TRANSBIG | | 197 (25%) | 30 (21%) | 153 (31%) | 14 (8.6%) |
| UNT | | 124 (16%) | 47 (33%) | 68 (14%) | 9 (5.6%) |

[1]Mean (SD); n (%)

**Table S4:** Relapse Free Survival - Patient Characteristics

| | | | Clock Function | | |
|---|---|---|---|---|---|
| **Variable** | **N** | **Overall**, N = 764[1] | **Good**, N = 184[1] | **Average**, N = 495[1] | **Bad**, N = 85[1] |
| **Theta** | 764 | 0.25 (0.18) | 0.06 (0.02) | 0.26 (0.10) | 0.61 (0.13) |
| **Age** | 764 | 52.53 (12.57) | 53.36 (12.70) | 52.49 (12.77) | 51.01 (10.99) |
| **Treatment** | 764 | | | | |
| No Treatment | | 478 (63%) | 122 (66%) | 316 (64%) | 40 (47%) |
| Treatment | | 286 (37%) | 62 (34%) | 179 (36%) | 45 (53%) |
| **Tumour Size** | 763 | | | | |
| T1 | | 295 (39%) | 73 (40%) | 194 (39%) | 28 (33%) |
| T2 | | 365 (48%) | 82 (45%) | 247 (50%) | 36 (42%) |
| T3_4 | | 103 (13%) | 28 (15%) | 54 (11%) | 21 (25%) |
| (Missing) | | 1 | 1 | 0 | 0 |
| **ER Status** | 754 | | | | |
| 0 | | 210 (28%) | 48 (27%) | 141 (29%) | 21 (25%) |
| 1 | | 544 (72%) | 132 (73%) | 349 (71%) | 63 (75%) |
| (Missing) | | 10 | 4 | 5 | 1 |
| **Grade** | 738 | | | | |
| 1 | | 137 (19%) | 29 (17%) | 95 (20%) | 13 (15%) |
| 2 | | 329 (45%) | 86 (50%) | 204 (42%) | 39 (46%) |
| 3 | | 272 (37%) | 56 (33%) | 184 (38%) | 32 (38%) |
| (Missing) | | 26 | 13 | 12 | 1 |
| **PAM50** | 764 | | | | |
| Basals | | 118 (15%) | 26 (14%) | 84 (17%) | 8 (9.4%) |
| HER2s | | 96 (13%) | 21 (11%) | 65 (13%) | 10 (12%) |
| LuminalA | | 351 (46%) | 97 (53%) | 219 (44%) | 35 (41%) |
| LuminalB | | 167 (22%) | 29 (16%) | 109 (22%) | 29 (34%) |
| Normals | | 32 (4.2%) | 11 (6.0%) | 18 (3.6%) | 3 (3.5%) |
| **Centre** | 764 | | | | |
| Rem_Cent_1 | | 103 (13%) | 22 (12%) | 64 (13%) | 17 (20%) |
| Rem_Cent_2 | | 58 (7.6%) | 13 (7.1%) | 35 (7.1%) | 10 (12%) |
| Rem_Cent_3 | | 17 (2.2%) | 4 (2.2%) | 10 (2.0%) | 3 (3.5%) |
| Rem_Cent_4 | | 32 (4.2%) | 9 (4.9%) | 13 (2.6%) | 10 (12%) |
| TRANSBIG | | 197 (26%) | 30 (16%) | 153 (31%) | 14 (16%) |
| UNT | | 124 (16%) | 47 (26%) | 68 (14%) | 9 (11%) |
| UPP | | 233 (30%) | 59 (32%) | 152 (31%) | 22 (26%) |

[1]Mean (SD); n (%)

**Table S5:** Model tests for the OS group (Remagus: IHC)

| | Model Tests | Discrimination Indexes |
|---|---|---|
| Obs 216 | LR $\chi^2$ 33.10 | $R^2$ 0.164 |
| Events 43 | d.f. 6 | $R^2_{6,216}$ 0.118 |
| Center -0.8164 | Pr($> \chi^2$) 0.0000 | $R^2_{6,43}$ 0.468 |
| | Score $\chi^2$ 34.59 | $D_{xy}$ 0.510 |
| | Pr($> \chi^2$) 0.0000 | |

**Table S6:** Likelihood Ratio Statistics for the OS group (Remagus: IHC)

| | $\chi^2$ | d.f. | $P$ |
|---|---|---|---|
| Theta | 7.41 | 2 | 0.0247 |
| Nonlinear | 7.40 | 1 | 0.0065 |
| size | 14.81 | 1 | 0.0001 |
| er | 0.01 | 1 | 0.9114 |
| pgr | 6.23 | 1 | 0.0126 |
| her2 | 2.49 | 1 | 0.1143 |
| **TOTAL** | 33.10 | 6 | <0.0001 |

**Table S7:** Resampling Validation for the OS group (Remagus: IHC)

| Index | Original Sample | Training Sample | Test Sample | Optimism | Corrected Index | Successful Resamples |
|---|---|---|---|---|---|---|
| $D_{xy}$ | 0.51 | 0.54 | 0.48 | 0.06 | 0.45 | 300 |
| Slope | 1.00 | 1.00 | 0.83 | 0.17 | 0.83 | 300 |

**Table S8:** Model tests for the OS group

| | Model Tests | Discrimination Indexes |
|---|---|---|
| Obs 422 | LR $\chi^2$ 32.63 | $R^2$ 0.088 |
| Events 92 | d.f. 6 | $R^2_{6,422}$ 0.061 |
| Centre -2.1202 | Pr(> $\chi^2$) 0.0000 | $R^2_{6,92}$ 0.251 |
| | Score $\chi^2$ 35.80 | $D_{xy}$ 0.292 |
| | Pr(> $\chi^2$) 0.0000 | |

**Table S9:** Likelihood Ratio Statistics for the OS group

| | $\chi^2$ | d.f. | $P$ |
|---|---|---|---|
| Theta | 11.36 | 2 | 0.0034 |
| Nonlinear | 11.06 | 1 | 0.0009 |
| treatment | 6.85 | 1 | 0.0088 |
| size | 15.16 | 1 | 0.0001 |
| age | 5.11 | 2 | 0.0775 |
| Nonlinear | 3.62 | 1 | 0.0570 |
| **TOTAL** | 32.63 | 6 | <0.0001 |

**Table S10:** Resampling Validation for the OS group

| Index | Original Sample | Training Sample | Test Sample | Optimism | Corrected Index | Successful Resamples |
|---|---|---|---|---|---|---|
| $D_{xy}$ | 0.45 | 0.47 | 0.42 | 0.05 | 0.39 | 300 |
| Slope | 1.00 | 1.00 | 0.86 | 0.14 | 0.86 | 300 |

**Table S11:** Model tests for the DMFS group

| | Model Tests | Discrimination Indexes |
|---|---|---|
| Obs 522 | LR $\chi^2$ 11.86 | $R^2$ 0.026 |
| Events 117 | d.f. 5 | $R^2_{5,522}$ 0.013 |
| Centre -0.6505 | Pr(> $\chi^2$) 0.0367 | $R^2_{5,117}$ 0.057 |
| | Score $\chi^2$ 18.31 | $D_{xy}$ 0.111 |
| | Pr(> $\chi^2$) 0.0026 | |

**Table S12:** Likelihood Ratio Statistics for the DMFS group

| | $\chi^2$ | d.f. | $P$ |
|---|---|---|---|
| Theta | 6.64 | 2 | 0.0361 |
| Nonlinear | 2.11 | 1 | 0.1468 |
| size | 4.80 | 1 | 0.0285 |
| age | 1.40 | 2 | 0.4978 |
| Nonlinear | 0.43 | 1 | 0.5104 |
| **TOTAL** | 11.86 | 5 | 0.0367 |

**Table S13:** Resampling Validation for the DMFS group

| Index | Original Sample | Training Sample | Test Sample | Optimism | Corrected Index | Successful Resamples |
|---|---|---|---|---|---|---|
| $D_{xy}$ | 0.35 | 0.37 | 0.32 | 0.05 | 0.30 | 296 |
| Slope | 1.00 | 1.00 | 0.72 | 0.28 | 0.72 | 296 |

**Table S14:** Model tests for the RFS group

| | Model Tests | Discrimination Indexes |
|---|---|---|
| Obs 775 | LR $\chi^2$ 23.55 | $R^2$ 0.032 |
| Events 236 | d.f. 8 | $R^2_{8,752}$ 0.02 |
| Centre -0.8868 | Pr($>\chi^2$) 0.0027 | $R^2_{8,226}$ 0.064 |
| | Score $\chi^2$ 28.67 | $D_{xy}$ 0.175 |
| | Pr($>\chi^2$) 0.0004 | |

**Table S15:** Likelihood Ratio Statistics for the RFS group

| | $\chi^2$ | d.f. | $P$ |
|---|---|---|---|
| Theta (Factor+Higher Order Factors) | 13.08 | 4 | 0.0109 |
| All Interactions | 9.60 | 2 | 0.0082 |
| Nonlinear (Factor+Higher Order Factors) | 10.32 | 2 | 0.0057 |
| size (Factor+Higher Order Factors) | 16.84 | 3 | 0.0008 |
| All Interactions | 9.60 | 2 | 0.0082 |
| age | 1.35 | 2 | 0.5101 |
| Nonlinear | 1.29 | 1 | 0.2563 |
| treatment | 4.48 | 1 | 0.0344 |
| **TOTAL** | 23.55 | 8 | 0.0027 |

**Table S16:** Resampling Validation for the RFS group

| Index | Original Sample | Training Sample | Test Sample | Optimism | Corrected Index | Successful Resamples |
|---|---|---|---|---|---|---|
| $D_{xy}$ | 0.28 | 0.30 | 0.25 | 0.05 | 0.23 | 300 |
| Slope | 1.00 | 1.00 | 0.76 | 0.24 | 0.76 | 300 |

**Table S17:** Probes used by TimeTeller analysis. Probe ID to Gene symbol conversion

| ID | Gene symbol |
|---|---|
| *202861_at* | *PER1* |
| *205251_at* | *PER2* |
| *209750_at* | *NR1D2* |
| *209782_s_at* | *DBP* |
| *209824_s_at* | *ARNTL* |
| *210971_s_at* | *ARNTL* |
| *213462_at* | *NPAS2* |
| *221045_s_at* | *PER3* |
| *39549_at* | *NPAS2* |
| *205459_s_at* | *NPAS2* |

### Part III

### Supplementary Figures

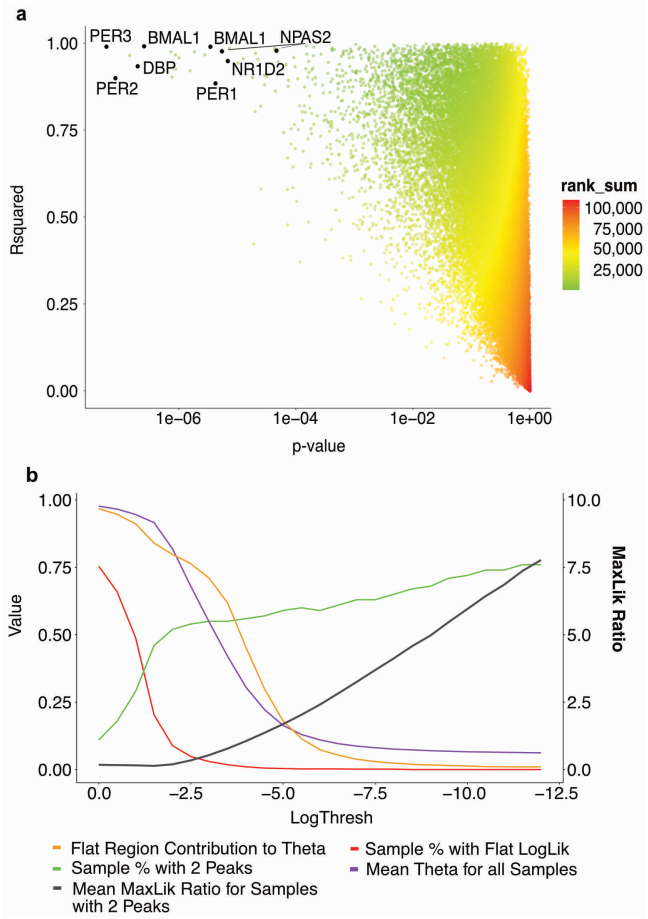


**Figure S1: Preparation of the training model. a.** Probes and genes to include in the REV are chosen in terms of their rhythmicity quality and syncronisation across tissues and individuals. This analysis, which is detailed in Fig B in S1 Appendix of (*23*), is important to choose a panel of genes with good circadian rhythmicity combined with minimal variation across the relevant tissues or individuals and to try to ensure it provides a faithful representation. Population cosinor $p$-value (rhythmicity detection) and $R^2$ (proportion of the variance explained by the rhythm) thresholds are used to pick the most rhythmic and synchronous genes. The figure is a scatter plot of these quantities for all genes using the training data. The $R^2$ is calculated across individuals. **b.** Summary stats for the metrics used to choose the log threshold selection. As discussed in Section S4, we want to reveal the relevant structure (i.e. % with 2 peaks and flat region contribution), while not deflating $\Theta$ too much. The region between -3.5 and -5 provides the best balance, so we choose Logthresh = -4.

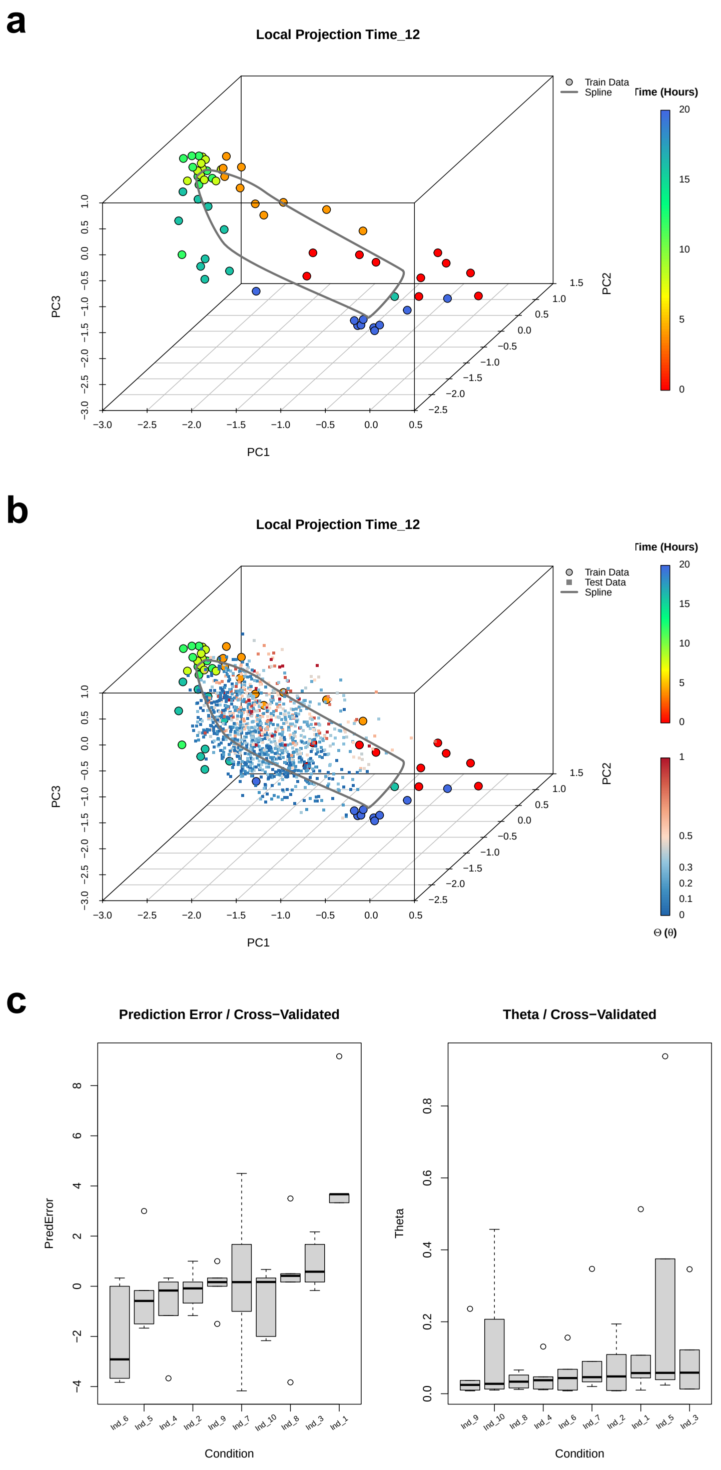


**Figure S2: Visualisation of the training model. a.** Visualisation of one of the local projections for the training model. Local projection time chosen to be $T = 12h$. Training samples displayed as circles and are colored by the known collection time. Projection is onto principal components obtained as described in (*23*). Grey line is the interpolated periodic spline. **b.** Leave-one-individual-out cross-validation of the TimeTeller model. Individuals with systematic prediction error (*Ind_6* and *Ind_1*) are consistent across the normalisation methods (Left). None of the individuals display evidence of perturbed clock (Right). **c.** Same as **a** but with the test data projected (small squares). Darker color corresponds to higher $\Theta$. See (*23*) for a detailed analysis of this dataset.

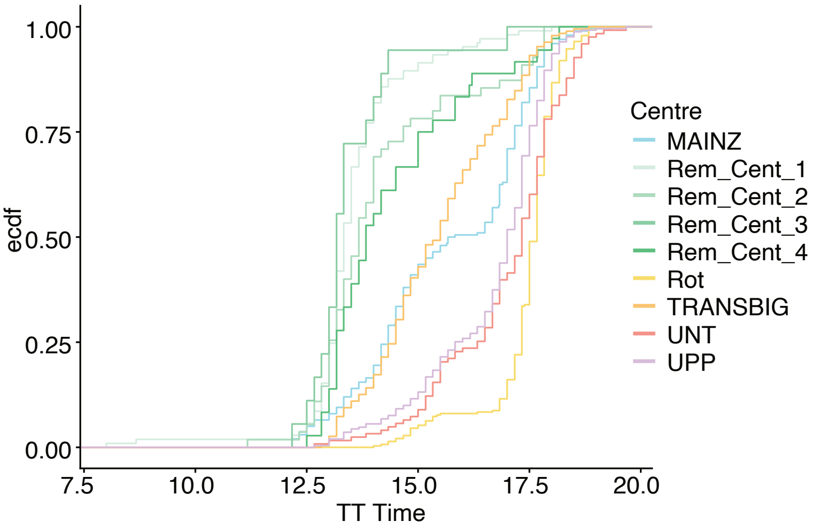


**Figure S3: TT Timing distributions by centre.** ECDF curves corresponding to TT timing per centre. Separation of Remagus centres earlier in the day is apparent.

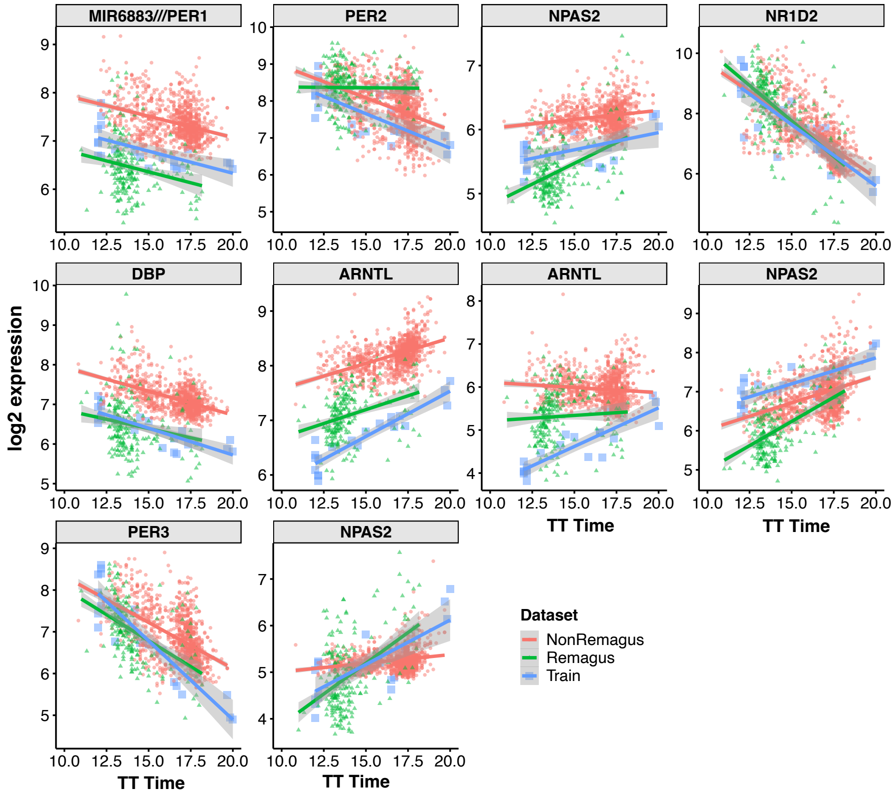


**Figure S4:** $\log_2$ **expression levels against TT time.** $\log_2$ expression of the probes corresponding to genes in the REV. In this figure the training data is given its TT time rather than its sample time. There is a definite temporal trend during the daytime.

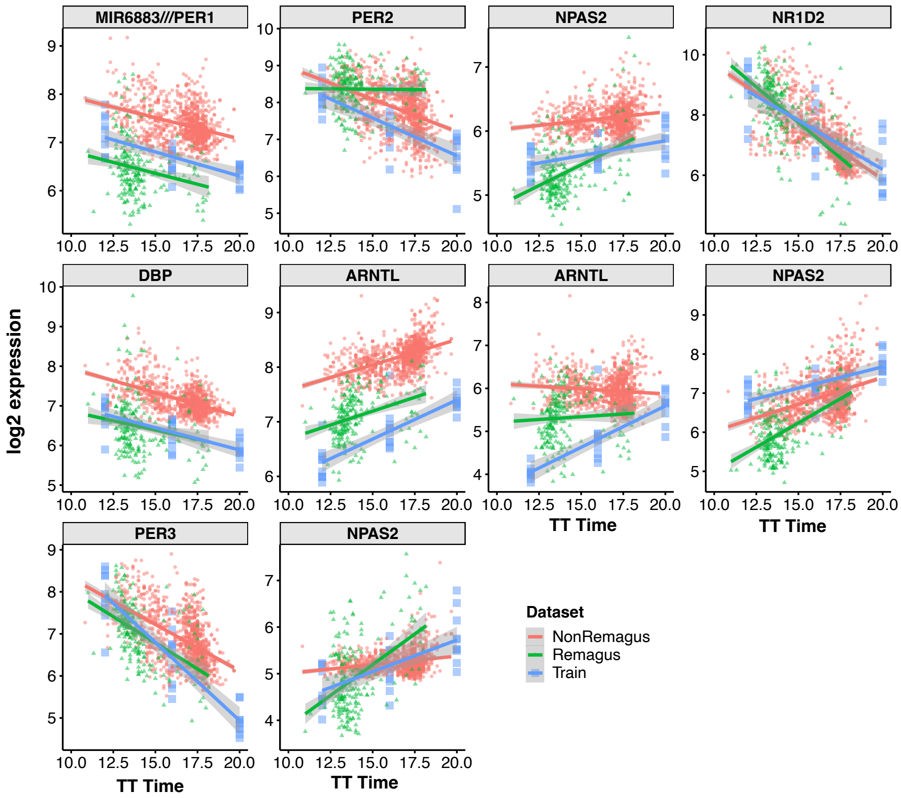


**Figure S5:** $\log_2$ **expression levels against TT time.** $\log_2$ expression of the probes corresponding to genes in the REV. In this figure the training data is given its sample time and not its TT time. There is a definite temporal trend during the daytime.

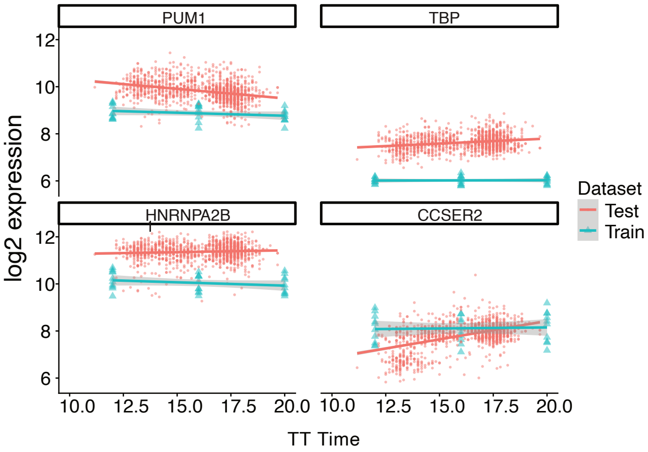


**Figure S6: Housekeeper gene levels against TT time.** Housekeeper gene $\log_2$ expression of some of the housekeeper genes commonly used in breast cancer research. As expected and contrary to the clock genes shown in the main text, there is no strong temporal trend.

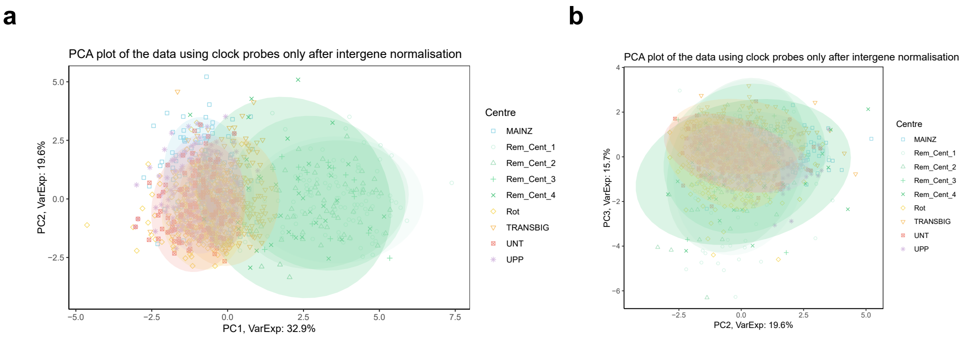


**Figure S7: PCA plots of clock probes used in the analysis. a**. PCA plot (PC1/PC2) showing test data projections. Only clock probes were used and SVD was performed on intergene normalised data. Visual appearance of batch effect for the Remagus centres could be explained by the TT timing differences. **b**. Same as **a** but for PC2/PC3. No batch effect apparent.

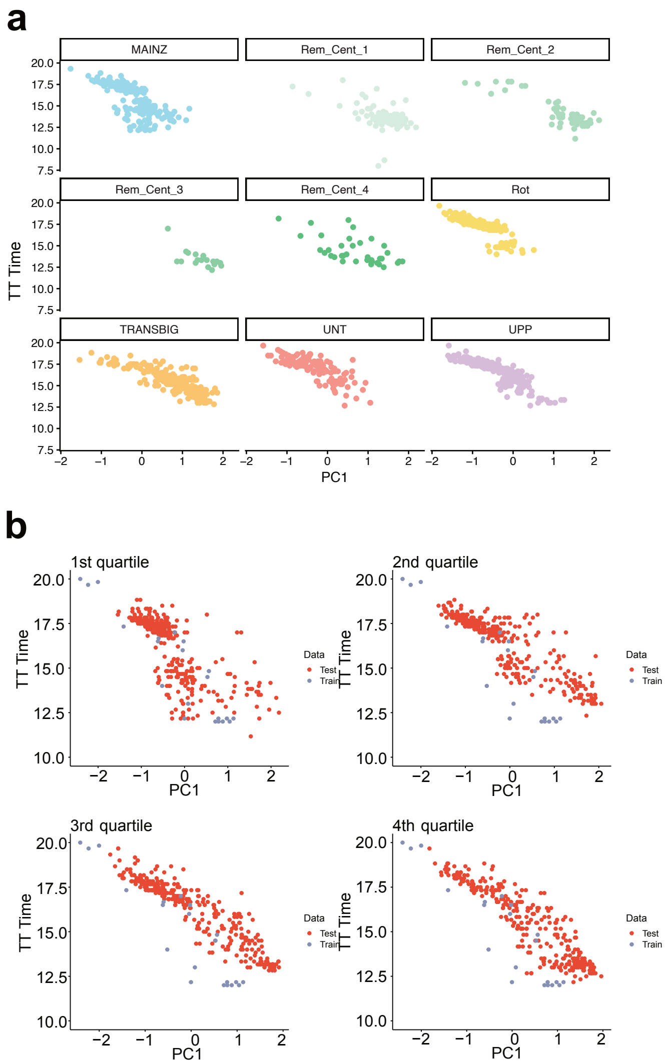


**Figure S8: Relationship between TT Time and projection onto first principal component of the REVs. a.** Projection of the test data onto test principal components (*PC1*). Faceted by centre. Most centres display spread throughout the day. **b.** Same as in Fig. 1F but split by *ML* quartiles. Higher spread is apparent for the bottom two quartiles (top panels) compared to the top two quartiles (bottom panels).

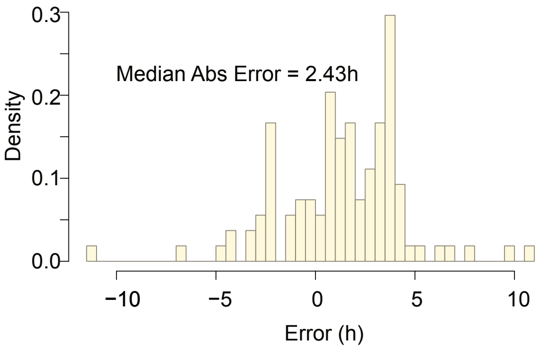


**Figure S9:** Histogram of deviations of the sample time , when known, from the TT time (*N* = *108*). Even though all the samples are predicted to come from the day time, median absolute error is high at 2.43h.

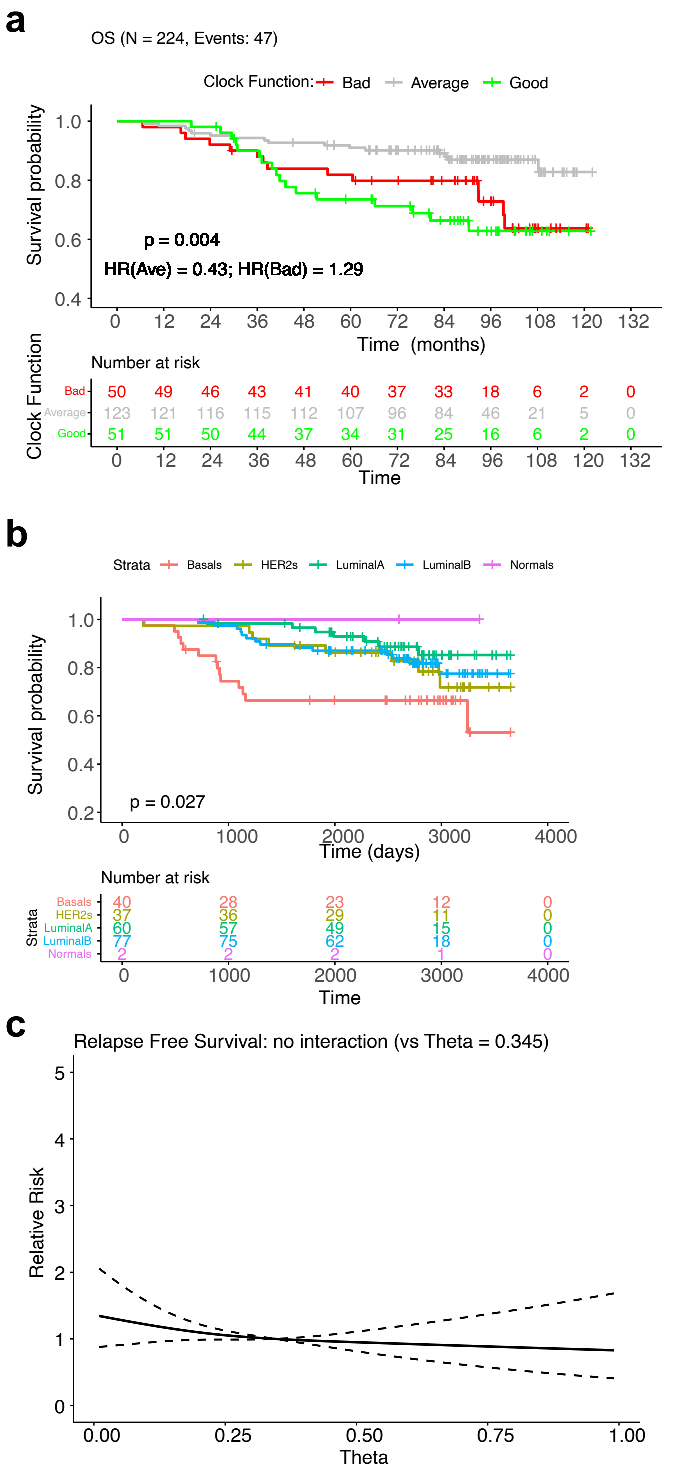


**Figure S10: Kaplan-Meier survival plots. a.** Kaplan-Meier survival plot showing differences in overall survival (OS) for patients with the lowest 20% of Θs (green), the highest 20% of Θs (red) and the middle 60% of Θs (grey). **b.** Kaplan-Meier survival plot showing differences in overall survival (OS) for patients with different predicted PAM50 subtypes. **c.** Relative risk as a function of Θ for the patients with available RFS information from the multivariate Cox model.

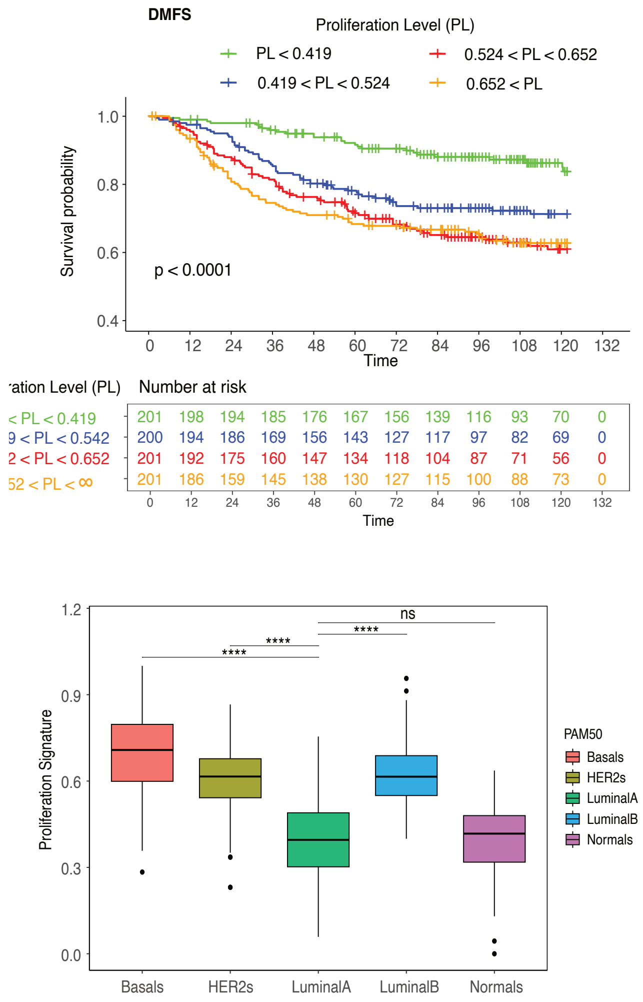


**Figure S11: Proliferation score. a.** Kaplan-Meier survival plot showing differences in distant metastasis free survival (DMFS) for patients with different inferred proliferation score. **b.** Boxplot showing differences in absolute value of *ssGSEA* proliferation signature grouped by inferred PAM50 subtype. As expected, *Basals* display the highest score, whereas *Luminal A* the lowest.

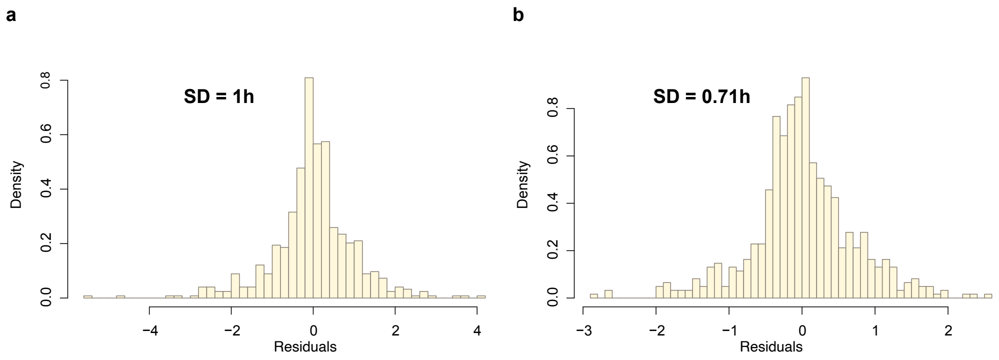


**Figure S12: a, b**. Histogram estimating $P(T|g_1)$ (as in the main text) for (a) the bottom 50% of ML values and (b) the top 50% of ML values. These are calculated by fitting a cubic spline to the corresponding scatter plot of TT time against $g_1$ and then calculating the horis=zonatl distance of each data point from the spline. As seen in Fig. S8b, the spread is lower for higher ML samples (SD = 0.71h vs SD = 1h).

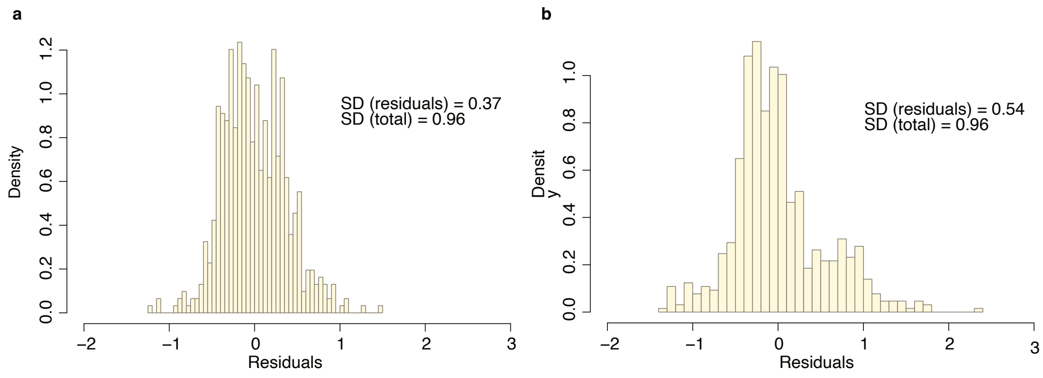


**Figure S13: Histogram estimating $P(g_1|T)$ (as in the main text). a, b**. Histogram estimating $P(g_1|T)$ (as in the main text). This is done by fitting cubic spline which is an estimate of the mean of the projection $g_1$ on to the first PC as a function of the TT time $T$. a. For the top 50% of ML values. (b) For the bottom 50% of ML values (b). As seen in Fig. S8b, the spread is lower for higher ML samples (SD = 0.37 vs SD = 0,54). The spread is defined to be the extent of the central 90% of the data i.e. after deleting the top and bottom 5%

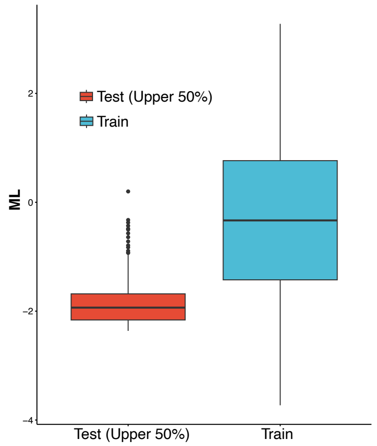


**Figure S14: Distributions of the values of the Maximum Likelihood (ML).** Density plot showing distribution of the test data (top 50% of ML values) and cross-validated train data.

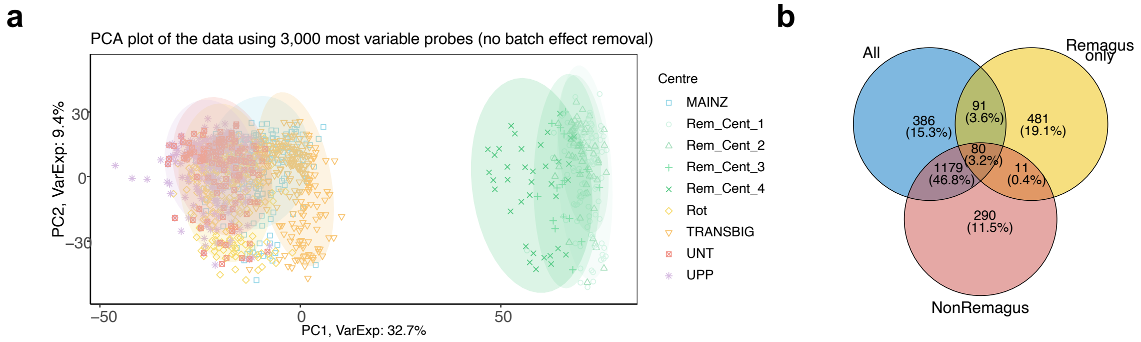


**Figure S15: PCA plots of most variable genes. a.** PCA plot (PC1/PC2) showing test data projections using 3,000 most variable genes. Batch effect evident for the Remagus centres. **b**. Venn diagram summarising DE analysis results for the combined test data (*blue*), Remagus only (*yellow*) and NonRemagus data (*red*). Overlap between the Remagus and NonRemagus is statistically significant (using permutation test) but limited in size potentially due to different tumor characteristics.

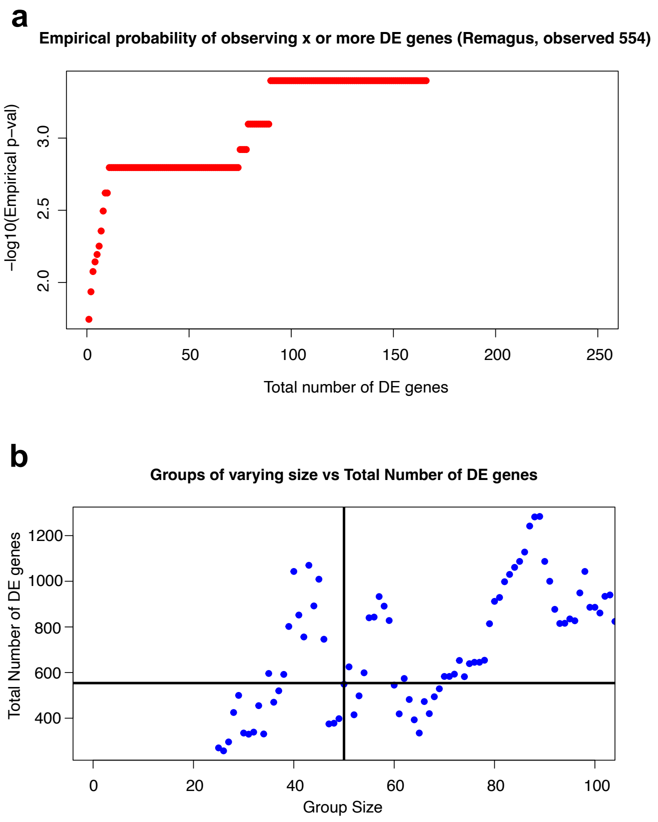


**Figure S16: Using the Θ stratification to find differentially expressed genes. a**. The probability of observing the indicated number of DEGs for the permuted GCG and BCG groups for Remagus. **b.** The number of differentially expressed genes as a function of the size of the GCG and BCG.

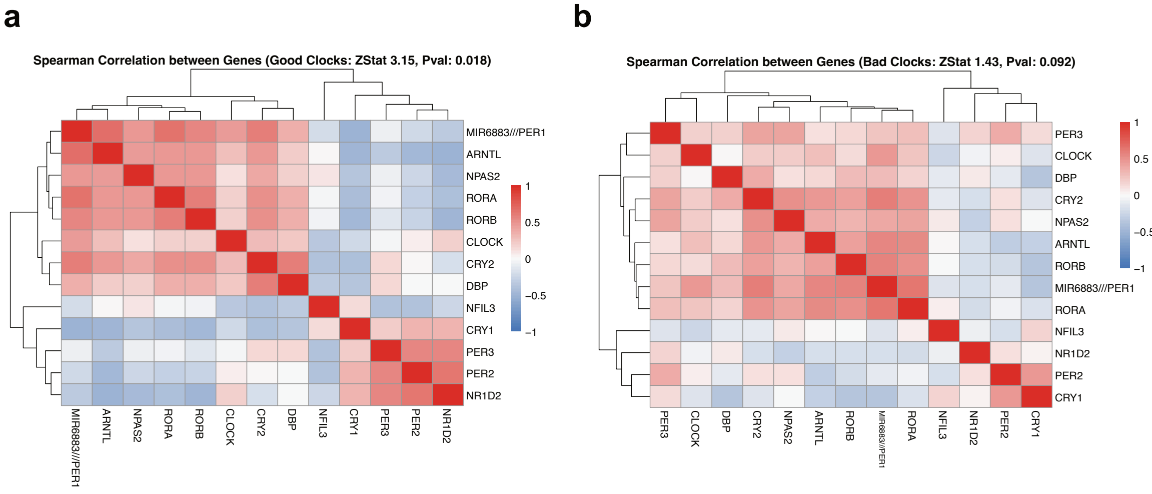


**Figure S17:** Using nCV (*24*) to test whether there is a functional clock network in population scale data using the reference training clock population (*23*). This is done using a combination of permutation testing and the Mantel test, which is a statistical test of the correlation between two matrices. The null hypothesis is that there is *no* relation between the two matrices so p-value below the threshold indicates that there is evidence to reject the null, hence the two correlation matrices are related (so null rejection implies working clock given the functioning clock in the reference data). In our case, the p-value for the TimeTeller predicted good clock group is 0.018 (**a**), whereas the p-value for the bad clock group is 0.092 (**b**), indicating consistency between the two methods.

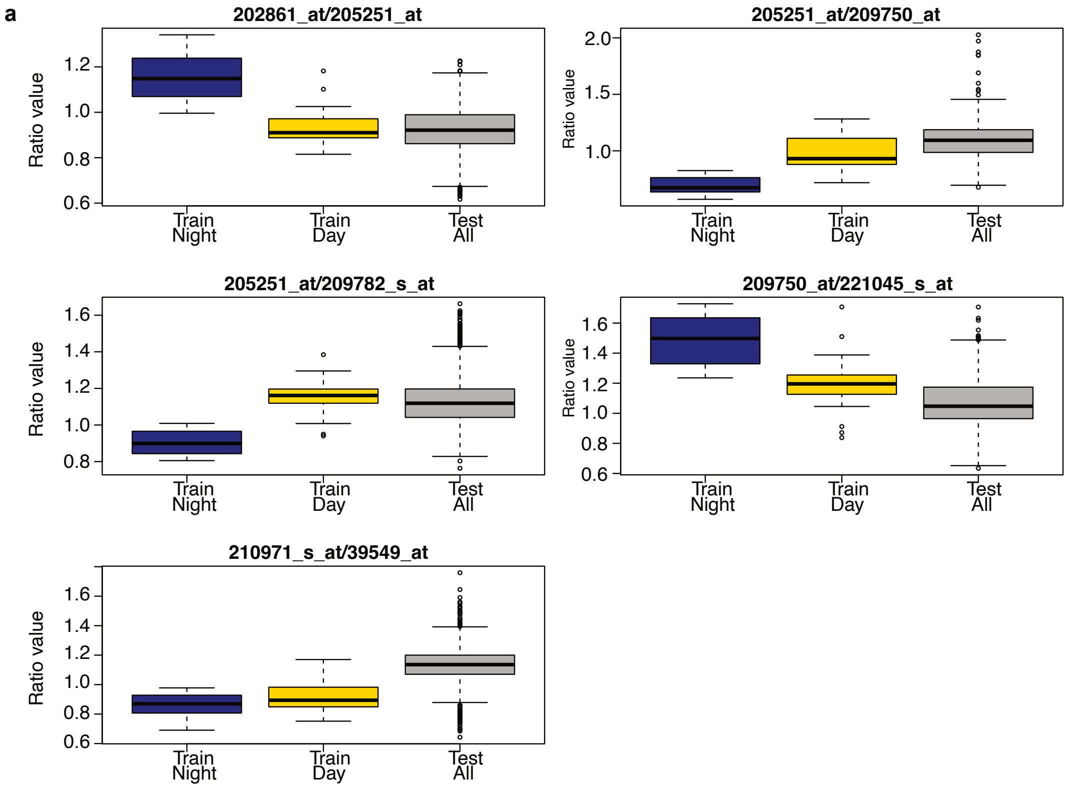


**Figure S18:** **a**. The five probe ratios used in the classifier. Box plots show performance on training and test samples.

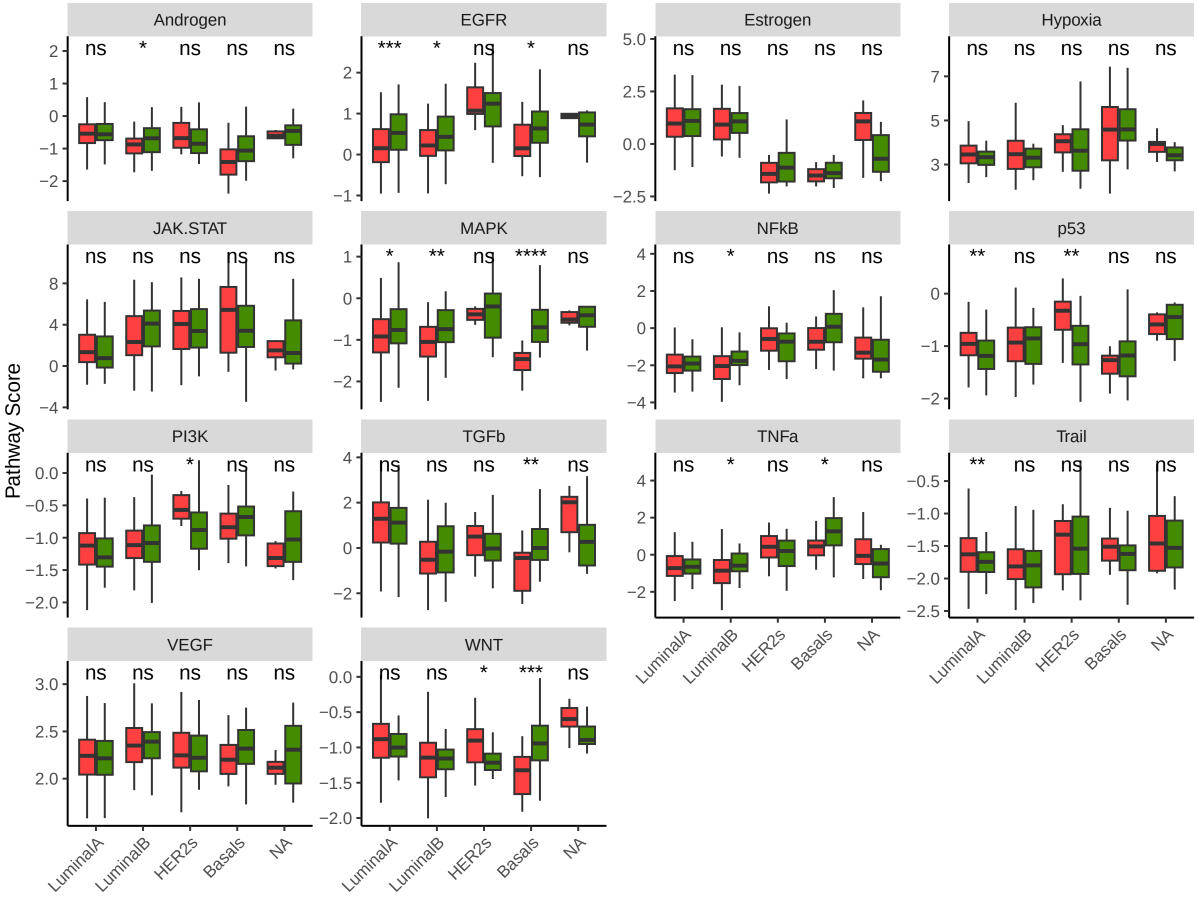


**Figure S19:** PROGENy-derived pathway activity scores for all 14 canonical pathways (Androgen, EGFR, Estrogen, Hypoxia, JAK-STAT, MAPK, NF$\kappa$B, p53, PI3K, TGF$\beta$, TNF$\alpha$, Trail, VEGF, WNT) across PAM50 breast cancer subtypes in the NonRemagus cohort. Samples are grouped by clock function: Good Clock (green) and Bad Clock (red) based on TimeTeller $\Theta$ quartiles. Boxplots show median (centre line), interquartile range (box), and 1.5× IQR (whiskers). Statistical significance assessed by Wilcoxon rank-sum test; significance markers indicate [$*P < 0.05$, $**P < 0.01$, $***P < 0.001$].

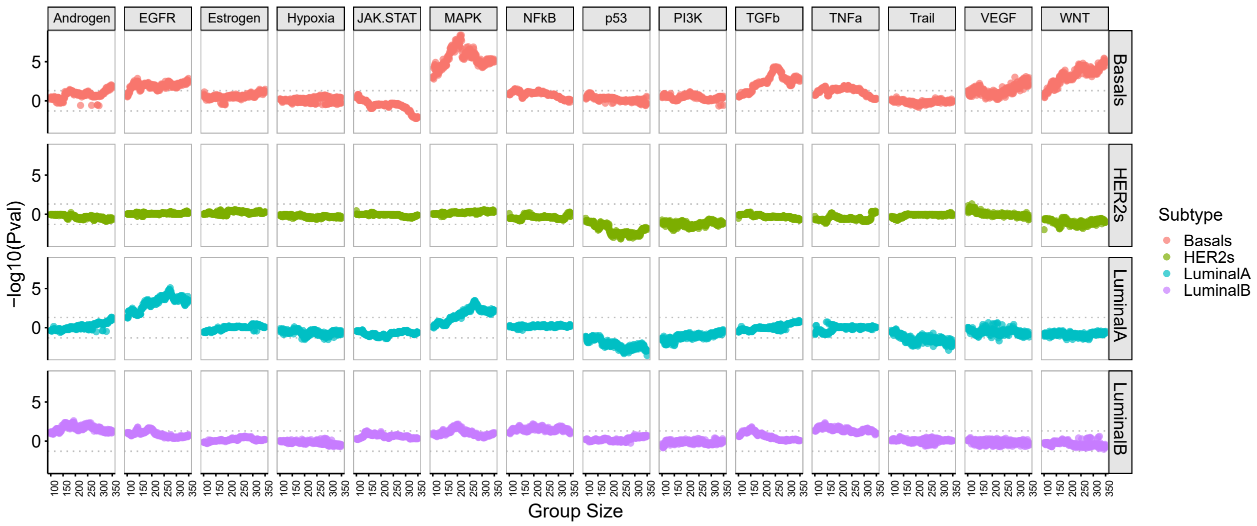


**Figure S20:** PROGENy-derived pathway activity scores for all 14 canonical pathways (Androgen, EGFR, Estrogen, Hypoxia, JAK-STAT, MAPK, NF$\kappa$B, p53, PI3K, TGF$\beta$, TNF$\alpha$, Trail, VEGF, WNT) across PAM50 breast cancer subtypes in the NonRemagus cohort. Samples are grouped by clock function and $-\log_{10}(P)$ from Wilcoxon rank-sum test calculated for varying Good/Bad Clock group sizes.

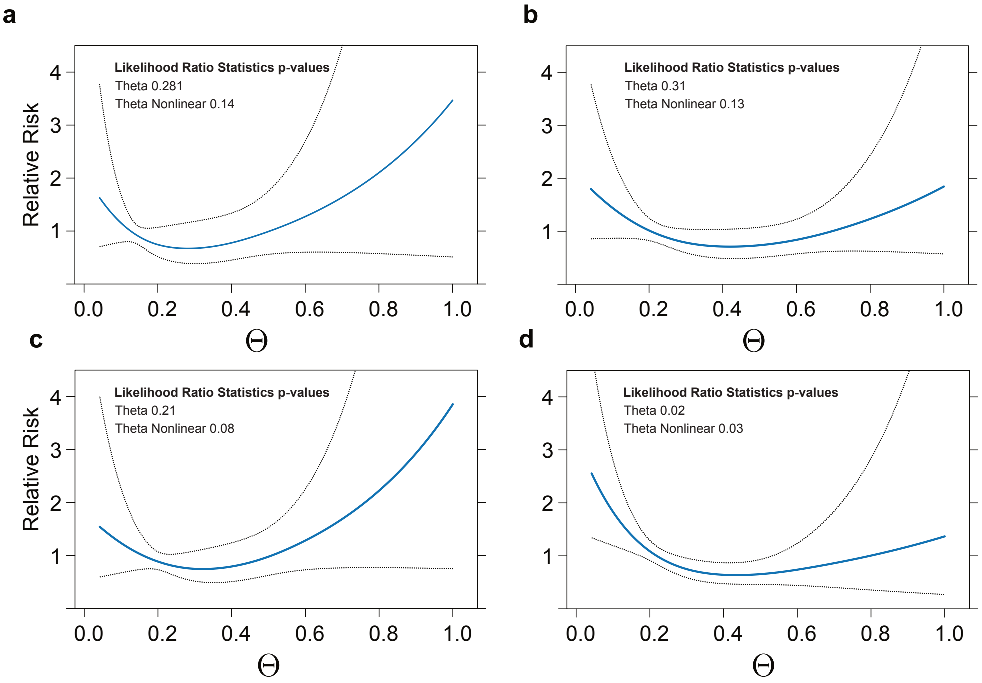


**Figure S21: Univariate survival analysis for OS by PAM50 subtype.** Relative risk as a function of $\Theta$ for four PAM50 subtypes (a) HER2, (b) Luminal A, (c) Luminal B, and (a) Basal.

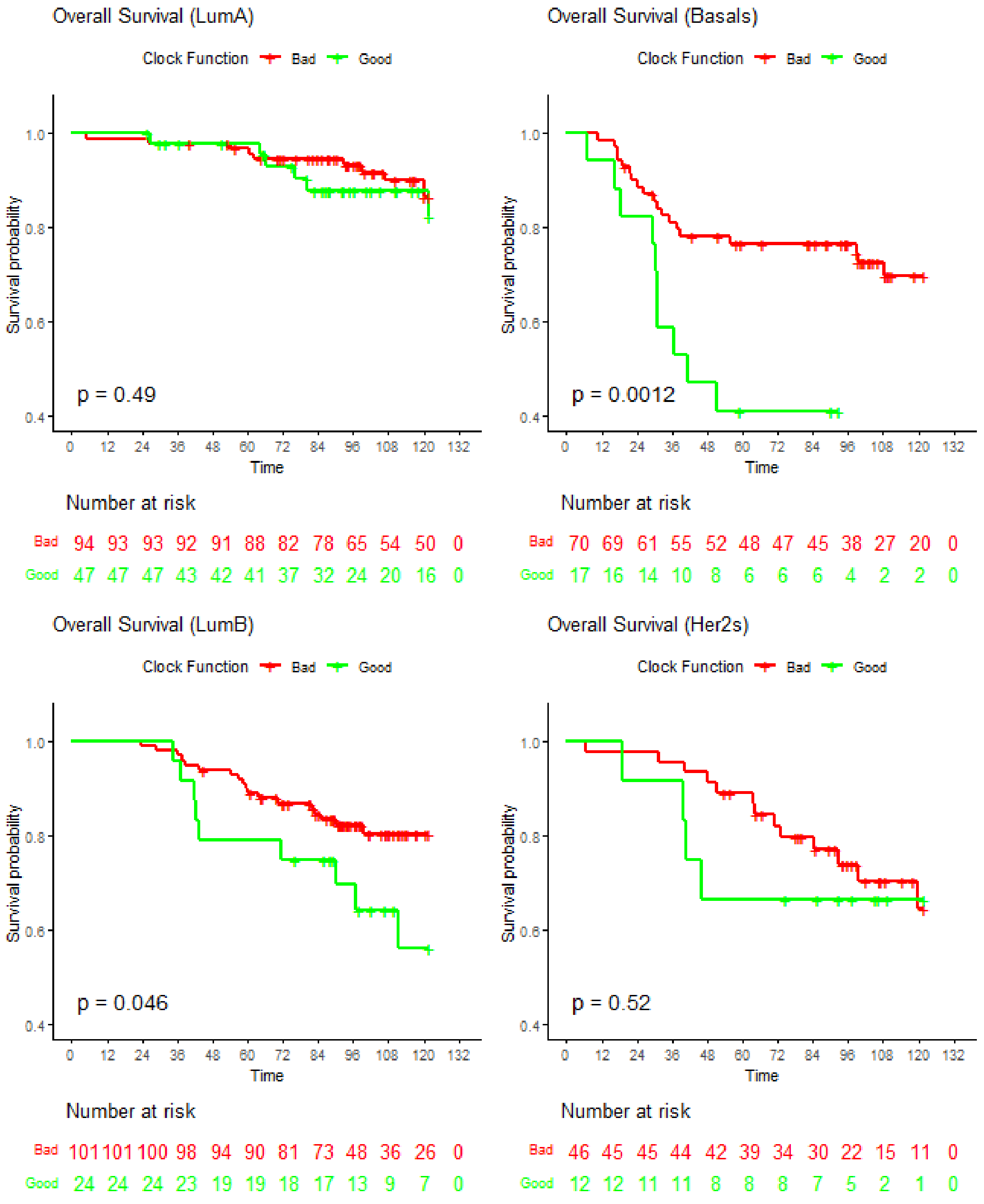


**Figure S22: Overall Survival KM curves by PAM50 subtype.** Analysis of the OS group. KM curves for the clock function split subgrouped by the PAM50 subtype. Logrank test and patient number at risk reported.

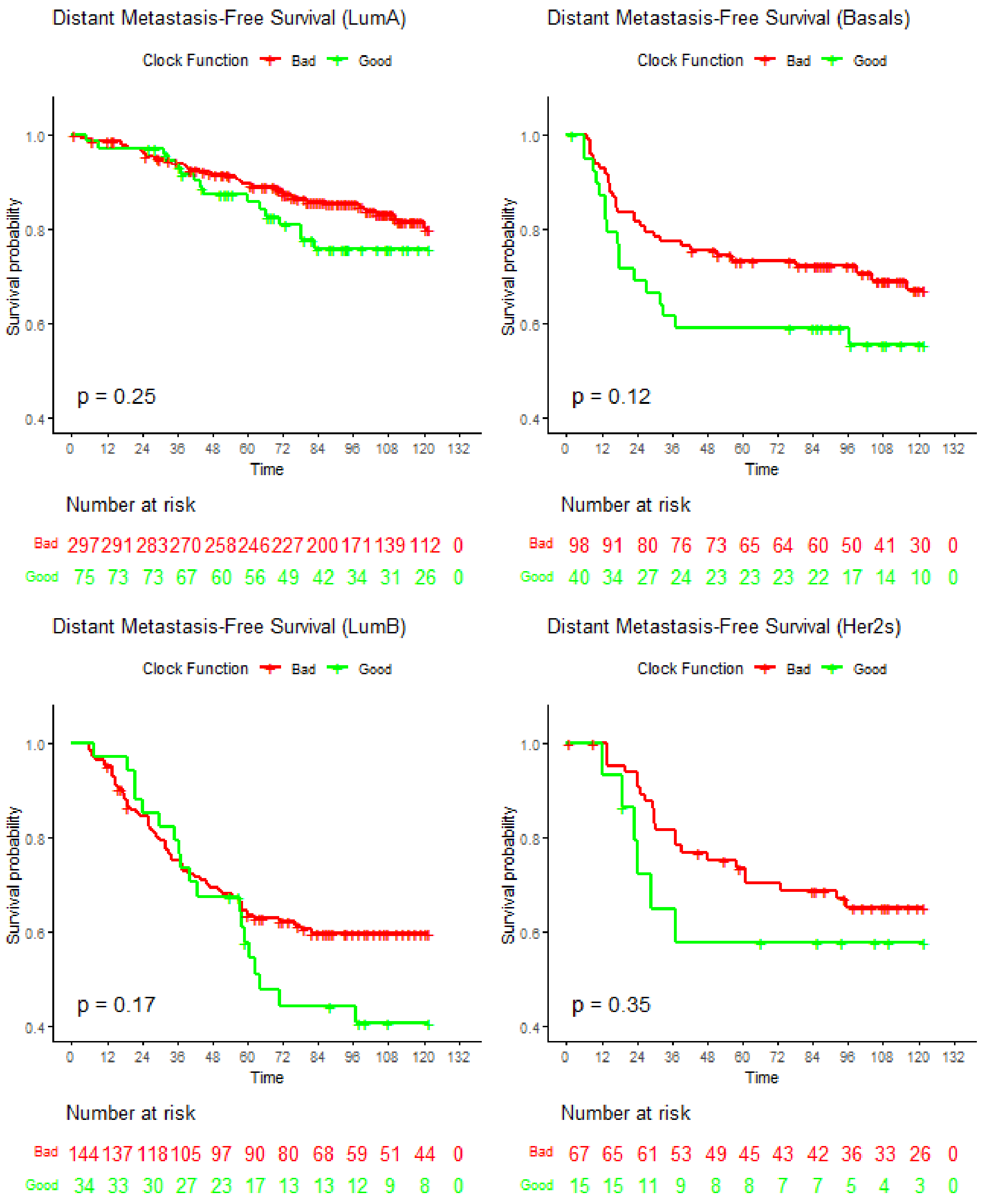


**Figure S23:** Analysis of the DMFS group. KM curves for the clock function split subgrouped by the PAM50 subtype. Logrank test and patient number at risk reported.

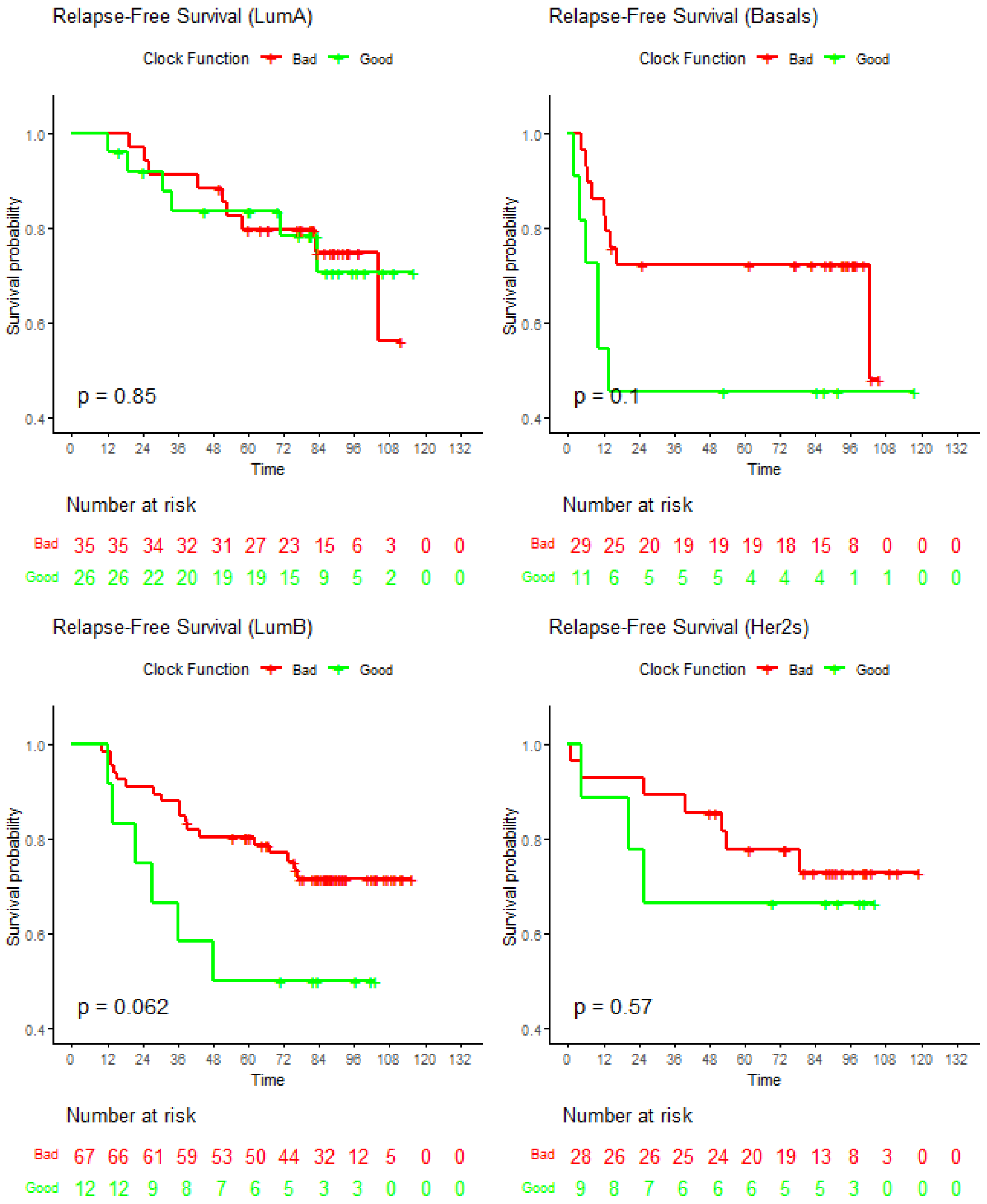


**Figure S24:** Analysis of the RFS group. KM curves for the clock function split subgrouped by the PAM50 subtype. Logrank test and patient number at risk reported.
